# Environmental identification of novel enzymes against heteroatomic plastics

**DOI:** 10.64898/2025.12.08.692966

**Authors:** Malthe Kjær Bendtsen, Andreas Møllebjerg, Samuel Peña-Díaz, Rosie Graham, Nicolai Claus Petersen, Bjørk Nolsøe Isaksen, Mathias Carstensen, Martin B. Johansen, Andreas Sommerfeldt, Allan R. Petersen, Iddi Khamisi Chuma, Cecilie Ryberg, Thomas Rea Wittenborn, Virginia Gichuru, Huabing Wang, Carsten Scavenius, Alexander Sandahl, Daniel E. Otzen

## Abstract

Better enzymes are needed to develop sustainable methods to recycle plastics with C-X heterobonds such as polyurethane (PUR) and nylon, for which no industrial-scale solutions exist. Current methods rely largely on sequence mining based on a small number of known enzymes. Here we expand the pool of PURases and nylonases by bioprospecting legacy plastic waste with fluorophore plastic mimics combined with FACS. We identify 29 plastic-degrading bacteria, from which 12 enzymes are identified by mass spectrometry and homology searches. Compared to existing enzymes, these enzymes are superior in thermostability and the ability to hydrolyse different high-molecular weight PUR oligomers and nylon textiles. To our knowledge, this is the first reported example of enzymes capable of hydrolysing longer chains of PUR and nylon. This study significantly increases the number of known PURases and nylonases and provides starting points for optimization campaigns through protein engineering and for *in silico* discovery.

## Introduction

Environmental plastic pollution has emerged, along with anthropogenic climate change, as one of the primary challenges of our time. To address this, efficient and economically competitive methods of recycling are sorely needed. Truly circular recycling requires the plastic to be deconstructed into monomeric building blocks which can be used to synthesize new polymers. Depolymerization by chemical catalysis requires high temperatures and high input purity, as blended plastics or additives can deactivate catalysts or produce toxic byproducts ^1-4^. Enzymes have recently emerged as attractive alternatives, or even partners, to chemical catalysts, due to their high selectivity and milder reaction conditions. While enzymatic recycling of polyethylene terephthalate (PET) waste shows promise at industrial scale^5^, enzymatic strategies are less developed for other heteroatomic plastics such as polyamides, or nylons, and polyurethane (PUR) although they represent promising targets for degradation. Nylons are predominately the polymers nylon 6 and nylon 6.6. PUR encompasses a range of carbamate-based polymers that include both thermoplastics (linear polymers) and thermosets (crosslinked polymers), often synthesized as copolymers consisting of an aromatic “hard segment” such as methylene diphenyl diisocyanate (MDI) or toluene diisocyanate (TDI) and a “soft segment” such as polyethylene or polypropylene glycol (PEG or PPG) or polycaprolactone (PCL)^6^. This necessitates selective degradation or extensive and costly downstream separation of monomers for recycling.

Most previously identified polyurethanases (PURases) have only been tested against ester-based polyurethane^6-13^ or have only documented activity against ethyl carbamate^14^, leaving to the best of our knowledge only ten PURases, proven to degrade the carbamate in PUR or PUR derived substrates: UMG-SP(1-3), *Nocardia farcinica* amidase, pQR3139, pQR3141, pQR3144, TflABH, MthABH and OspAmd^15-18^. They vary greatly in catalytic efficiency, and none have been shown to degrade untreated PUR. Additionally, their selectivities towards different formulations of PUR are largely undescribed. Likewise, many enzymes labelled as nylonases have only been tested on soluble amide compounds^19-21^, or biodegradable polyamides^22,23^, while those tested on recalcitrant nylon 6 or 6,6 show minimal activity^24,25^. Hence, discovery and characterisation of true PURases and nylonases are needed from which to optimize practical enzymatic recycling of PUR and nylon.

Existing discovery strategies of plastic-degrading enzymes include the repurposing of enzymes with activity against similar compounds^12,14^ or bioinformatic mining of protein databases for sequence- or structure-based homologues. Homology searches, however, only explore local sequence space, which limits options. Expansion to other regions of sequence space requires the discovery of novel non-homologous enzymes by functional screening, either by selective enrichment of microorganisms on plastic substrates (ability to metabolize plastic ^9,11,13,19,22,23^) or direct activity-based measurements (ability to cleave plastic bonds)^10,18^. Fluorescent probes are particularly useful, as they couple cleavage of the bond of interest to a fluorescent shift, allowing for highly sensitive detection of enzymatic activity. They have been used to discover enzymes such as hydrolases, lipases, nitroreductases and even PETases^26-28^. Utilising fluorescent probes for an activity-based screening allows isolation of enzymes from environmental samples, even those found in low abundance and with modest activity.

We present a new functional screening platform by which we discover twelve novel plastic-degrading enzymes, obtained from environmental microbiomes and exhibiting high structural and sequence diversity. Fluorescent probes targeting PURases and nylonases were used to isolate bacteria expressing these enzymes via fluorescence-assisted cell sorting (FACS). The corresponding enzymes were identified through sequential purification from the host bacteria, tracking their activity with the same probes and finally identifying them by mass spectrometry. The enzymes show diverse activity profiles across PUR and Nylon, with superior thermostability (melting temperature *t*_m_ up to 68.3 ± 0.1℃) and hydrolysis across more realistic substrates (high-MW PUR oligomers and Nylon textiles) than previously described enzymes.

### Development of a rapid functional screening method to isolate carbamate- and amide-cleaving bacterial strains

To expand the enzymatic toolbox for plastic depolymerization, we developed a rapid (>10^7^ bacterial cells/h) functional screening platform to screen bacteria containing putative PURases and nylonases from plastic-contaminated environments. Microorganisms are first extracted and incubated with fluorescent activity probes designed to mimic the chemical structure of the plastics (**Fig. 1a, Supplementary Figs. 1** and **2**). To our knowledge, this study is the first reported example of enzymes capable of hydrolysing longer chains of PUR and nylon and provides a platform for future optimization campaigns to develop enzymatic solutions to plastic degradation through *in silico* discovery and protein engineering). The PURase probe (PURp1) contains a carbamate bond between a bulky aromatic group and an aliphatic chain, while the nylonase probe (NYLp1) contains an amide bond between two aliphatic chains (**Supplementary Table S1**). Enzymatic cleavage of PURp1 exposes the free 4-amino group, red-shifting fluorescence emission, while cleavage of NYLp1 releases the quencher and thus activates fluorescence^29^. Both probes are small, neutral and relatively hydrophobic, allowing passive diffusion into microorganisms^30^. The cleaved PURp1 can also diffuse into the bacteria (**Fig. S1a**). The fluorophore released by cleavage of NYLp1 is positively charged and cannot diffuse freely (**Fig. S2**), thus trapping intracellularly cleaved NYLp1 inside the cell while extracellularly cleaved probe may be electrostatically attracted to the predominantly anionic cell membrane. This allows both probes to stain bacteria with different expression patterns. Microorganisms with increased fluorescence are isolated by FACS at rates of up to 10.000 events per second (**Fig. 1a, Fig. S1bcd**).

**Figure 1.**
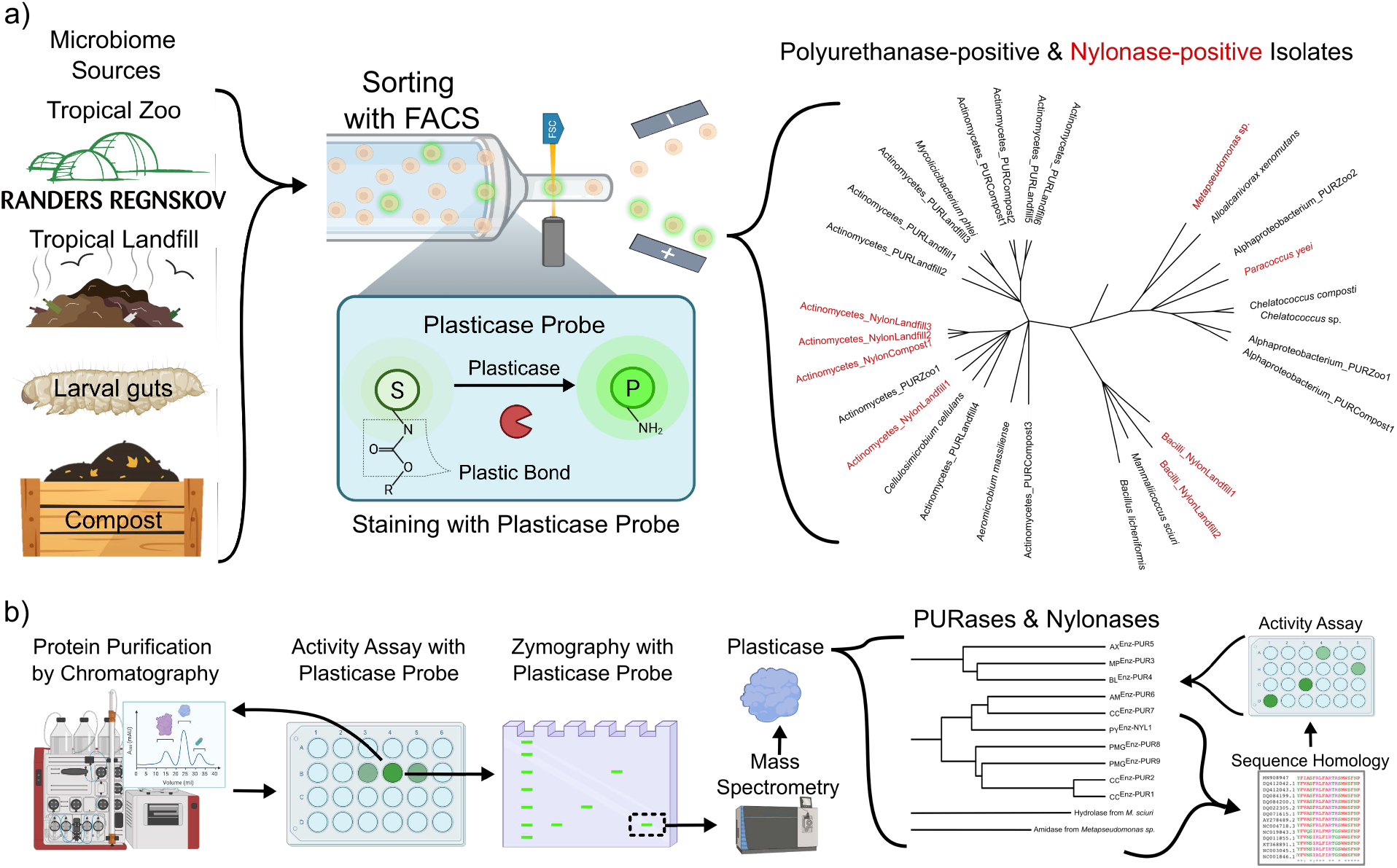
High-throughput functional screening platform to discover PURases and Nylonases (collectively termed plasticases). (a) Isolation of PURase- and Nylonase-positive bacteria from plastic-contaminated environments. Bacteria were extracted from plastic samples in a tropical zoo, a Kenyan landfill, a compost bin and larval guts. The bacteria were stained with a PURase or nylonase probe and sorted by FACS, based on the fluorescence signal of the cleaved probe. 21 PURase-positive and 8 nylonase-positive species were isolated. b) Identification of PURases and Nylonases from environmental isolates. The PURases and nylonases were purified from their isolated organisms, based on successive rounds of chromatography and activity assays with their respective probes. Purified enzyme were identified by mass spectrometry and expressed recombinantly. Additional enzymes were identified by searching the isolate genomes and public databases for homologues to the identified enzymes. In total, 10 PURases and 2 nylonases were identified.

To isolate microorganisms with plastic-depolymerizing enzymes, we collected plastic-waste samples (composition confirmed by ATR-FTIR (**Fig. S3**)) from humid and warm environments favouring bacterial growth and enzymatic thermal stability, exemplified by Kenyan landfills, a tropical zoo, PUR enriched compost^31,32^, and the guts of wax-worm reared on PUR foam^33^. From the landfill, FACS isolated eight PURase-positive species from polyurethane foam and wire and five nylonase-positive species from nylon nets. From the polyurethane in the tropical zoo, five PURase-positive species were isolated. From the compost, four PURases-positive species were isolated from the polyurethane foam and three from the elastane, while three nylonase-positive species were isolated from nylon fibers. From the wax-worm guts, one PURase-positive species was isolated. No bacterial species from polyethylene or polypropylene samples were consistently active against our probes.

In total, 21 PURase-positive and 8 nylonase-positive bacterial species were isolated, with relative activities spanning a >10.000-fold range (**Fig. S4**), highlighting the platform’s high sensitivity and broad dynamic range. The isolates were taxonomically diverse, comprising species from Actinomycetia α-proteobacteria, γ-proteobacteria and Bacilli. Despite this diversity, Actinobacteria were overrepresented for both plastic types. This may reflect the phylum’s broad enzymatic diversity, as indicated by their disproportionate contribution to degradation of xenobiotics^34,35^, as well as their high representation in the CAZy and PAZy databases^36,37^.

Although the platform preferentially enriches for cell-associated enzymes, of the 29 isolates, two nylonase-positive species secreted all their enzyme activity (**Fig. S4**). Detection of these species is probably facilitated by electrostatic attraction between the released fluorophore and the bacterial membrane.

### Identification and characterisation of putative PURases and nylonases

To identify active enzymes from these different species, the novel isolates were grown and lysed to extract the soluble fraction, followed by different combinations of non-denaturing liquid chromatography including anion exchange (AEX), size exclusion (SEC) and/or hydrophobic interaction (HIC), while monitoring PURase or nylonase (NYLp1) activity of individual fractions using fluorescent probes (**Fig. 1b** and **Fig. S5**). After iterative purification of the most active fractions, the potentially active enzymes were isolated (**Table S2**), fractions were digested, and proteins identified by mass spectrometry. Additionally, native-PAGE zymography aided the isolation of potentially active enzymes in *Alloalcanivorax xenomutans* and *Chelatococcus* sp (**Fig. 1b, Fig. S5** and **Table S2**).

We identified several enzymes active on the PURase probe. CC^EnZ-PUR1^ and CC^EnZ-PUR2^, from two different *Chelatococci* species, showed 89% sequence identity to each other but only 38% identity to the published PURase UMG-SP1^18^. BL^EnZ-PUR4^ from *Bacillus licheniformis* is a predicted *p*-nitrobenzyl esterase, a homolog of an esterase from *B. subtilis* identified as a PETase^43^. AX^EnZ-PUR5^ from *Alloalcanivorax xenomutans* has 36% identity to TfCa PETase from *Thermobifida fusca*. The actinobacterial enzymes AM^Enz-PUR6^ from *Aerobacterium massiliense* and CC^Enz-PUR7^ from *Cellulosimicrobium cellulans* were predicted as an acylamidase and a 6-aminohexanoate-cyclic-dimer hydrolase, respectively. In addition, we identified a metal-dependent hydrolase from *M. sciuri*, but the activity of the recombinant enzyme was too poor to merit further study on this probe.

As regards nylon degradation, MS-based screening identified a predicted (R)-stereoselective amidase from *Metapseudomonas* sp, but with insufficient activity levels in the recombinant form. We were unable to identify nylonases from isolates of the remaining seven nylonase-positive species. Instead, we used their genomes to perform a computational homology screening that comprises: *i)* the construction of an enzymatic library containing amidases, proteases, peptidases and hydrolases from our bacterial genomes; *ii)* sequence homology search among this library using plastic-degrading enzymes from this study and the literature *iii)* structural comparisons employing AlphaFold-derived models (**Fig. 1b**). In *P. yeii* this identified PY^EnZ-NYL1^, originally described as a 6-aminohexanoate-cyclic-dimer hydrolase, which had limited identity with NylA (37.34%) and CC^EnZ-PUR1^ (32.71%). Likewise, from the isolated *Mycolichibacterium phlei* genome we identified MP^EnZ-PUR3^, with 31.95% identify to AX^EnZ-PUR5^. Our success prompted a similar approach in a published PUR-derived metagenome^38^, leading to two pseudogenes coding for active enzymes, PMG^EnZ-PUR8^ and PMG^EnZ-PUR7 PUR9^ with 41 and 37% identity, respectively, to CC^EnZ-PUR1^.

In total this approach has now identified 10 novel PURases and 2 novel nylonases. Many of the enzymes are active on both PURp1 and NYLp1-3 **(Table 1** and **Fig. 3a)**. Use of multiple nylon fluorophores reduced false negatives caused by substrate specificity. Thus PY^EnZ-NYL1^, CC^EnZ-PUR2^ and PMG^EnZ-PUR9^ were much more active on NYLp2 than NYLp1 (**Fig. 3a** and **Fig. S7**). Activity on PURp1 varied widely, with CC^EnZ-PUR1^, CC^EnZ-PUR2^, AX^EnZ-PUR5^ and PMG^EnZ-PUR9^ having similar activity to UMG-SP3 while PMG^EnZ-PUR8^, PY^EnZ-NYL1^ and BL^EnZ-PUR4^ and AM^EnZ-PUR6^ had lower activities.

**Table 1.**

| Name | Bacterium of Origin | Isolated from | Enzyme class | Melting point | Activity on PURase probe (μmol min<sup>-1</sup> μmol<sup>-1</sup>) | Activity on DUE TDA | Activity on DUE-MDA | Activity on PUR oligomers | Activity on nylonase probe | Nylon dimers | Nylon 6,6 |
| --- | --- | --- | --- | --- | --- | --- | --- | --- | --- | --- | --- |
| **CC**<sup>EnZ-PUR1</sup> | *Chelatococcus composti* | PUR enriched compost | Amidase | 68.3 ± 0.1 | 4.0 ± 1.6 | + | + | + | + | + | + |
| **CC**<sup>EnZ-PUR2</sup> | *Chelatococcus* sp. | PUR enriched compost | Amidase | 58.3 ± 0.1 | 3.1 ± 0.6 | + | + | NA | + | + | + |
| **MP**<sup>EnZ-PUR3</sup> | *Mycolicibacterium phlei* | PUR enriched compost | Esterase | 50.4 ± 0.3 | 0.00222 ± 0.00004 | - | + | NA | - | NA | NA |
| **BL**<sup>EnZ-PUR4</sup> | *Bacillus licheniformis* | *Galleria mellonella* larvae gut | Esterase | 49.9 ± 0.3 | 0.0218 ± 0.0002 | - | + | NA | - | NA | NA |
| **AX**<sup>EnZ-PUR5</sup> | *Alloalcanivorax xenomutans* | Tropical zoo | Esterase | 49.7 ± 0.01 | 2.98 ± 0.02 | + | + | + | + | - | NA |
| **AM**<sup>EnZ-PUR6</sup> | *Aeromicrobium massiliense* | Tropical landfill | Amidase | 53.8 ± 0.1 | 0.00014 ± 0.00001 | + | + | + | + | - | NA |
| **CC**<sup>EnZ-PUR7</sup> | *Cellulosimicrobium cellulans* | Tropical landfill | Amidase | 64.1 ± 0.2 | 0.114 ± 0.003 | + | + | + | + | + | NA |
| **PMG**<sup>EnZ-PUR8</sup> | Plastic metagenome | Homology search | Amidase | 52.9 ± 0.1 | 0.67 ± 0.05 | + | + | NA | NA | NA | NA |
| **PMG**<sup>EnZ-PUR9</sup> | Plastic metagenome | Homology search | Amidase | 50.6 ± 0.1 | 13.4 ± 0.4 | + | + | + | + | + | NA |
| **PY**<sup>EnZ-NYL1</sup> | *Paracoccus yeii* | Nylon enriched compost | Amidase | 43.2 ± 0.1 | 0.44 ± 0.01 | - | + | NA | + | + | - |
| **NA** | *Metapseudomonas* sp. | Nylon enriched compost | (R)-stereoselective amidase | 62 ± 1 | NA | - | - | NA | + | NA | NA |
| **NA** | *Mammaliicoccus sciuri* | Tropical zoo | metal-dependent hydrolase | 58 ± 2 | NA | - | - | NA | + | NA | NA |
| **UMG-SP3** | Metagenome |  | Amidase |  | 21.0 ± 0.9 | + | + | NA | + | + | NA |

Using PURp1 for activity measurements, all enzymes except MP^EnZ-PUR3^ showed alkaline preference with activity optima around pH 8-11 (**Fig. S5b**). Optimum profiles were highly similar for enzymes within the same family. AX^EnZ-PUR5^ and BL^EnZ-PUR4^ showed a wide pH tolerance with >50% activity from pH 5-9 and optimum at pH 8.5. All amidases retained high activity at pH 8-9.5, some such as CC^EnZ-PUR2^ showed a broader pH range. The enzymes showed a range of thermostabilities with PY^EnZ-NYL1^ least stable (t_m_: 43.2 ± 0.1℃) and the most stable being CC^EnZ-PUR1^ (t_m_: 68.3 ± 0.1℃), CC^EnZ-PUR7^ (t_m_: 64 ± 0.2 ℃), and CC^EnZ-PUR2^ (t_m_: 58.3 ± 0.1℃) (**Fig. S5a** and **Table 1**), making them much more thermostable than published enzymes^18^.

As a first step to elucidate the enzymes’ ability to hydrolyse a substrate more representative of PUR, we synthesized two soluble dicarbamate compounds, namely diurethane ethylene (DUE)-MDA^39^ and the novel DUE-TDA (**Table S1** and **Supplementary Information**), representing MDI- and TDI-based PUR, respectively. These can potentially be degraded to mono-substituted monourethane ethylene (MUE)-MDA and MUE-TDA, and subsequently to 4,4-MDA and 2,4-TDA (cleavable bonds shown in **Table S1**). Enzyme activity was measured at 35°C (**Fig. 2ab**). Strikingly, all ten investigated enzymes except MP^EnZ-NYL2^ degraded DUE-MDA to at least MUE-MDA (**Fig. 2a**). For DUE-MDA, all PURases discovered successfully showed product release, highlighting the effective identification of enzymes capable of degrading the carbamate bond within soluble PUR oligomers. AX^EnZ-PUR5^, AM^Enz-PUR6^ and CC^Enz-PUR7^ released the highest amount of total product yield in 48h, with AM^Enz-PUR6^ and CC^Enz-PUR7^ showing full conversion of released MUE-MDA to 4,4-MDA with up to 40.9 ± 1.2 µg/ml and 47.1 ± 2.0 µg/ml 4-4 MDA detected respectively (**Fig. 2a**). In comparison, UMG-SP3 ^18^ only released 7.1 ± 4.4 µg/ml 4-4 MDA. Five enzymes (CC^EnZ-PUR1^, CC^EnZ-PUR2^, MP^EnZ-PUR3^, BL^EnZ-PUR4^ and PY^EnZ-NYL1^) degrade DUE-MDA to MUE-MDA only (**Fig. 2a**), while six (AX^EnZ-PUR5^, AM^Enz-PUR6^, CC^Enz-PUR7^, PMG^EnZ-PUR9^, PMG^EnZ-PUR8^ and UMG-SP3) process DUE-MDA via MUE-MDA to 4,4-MDA.

**Figure 2.**
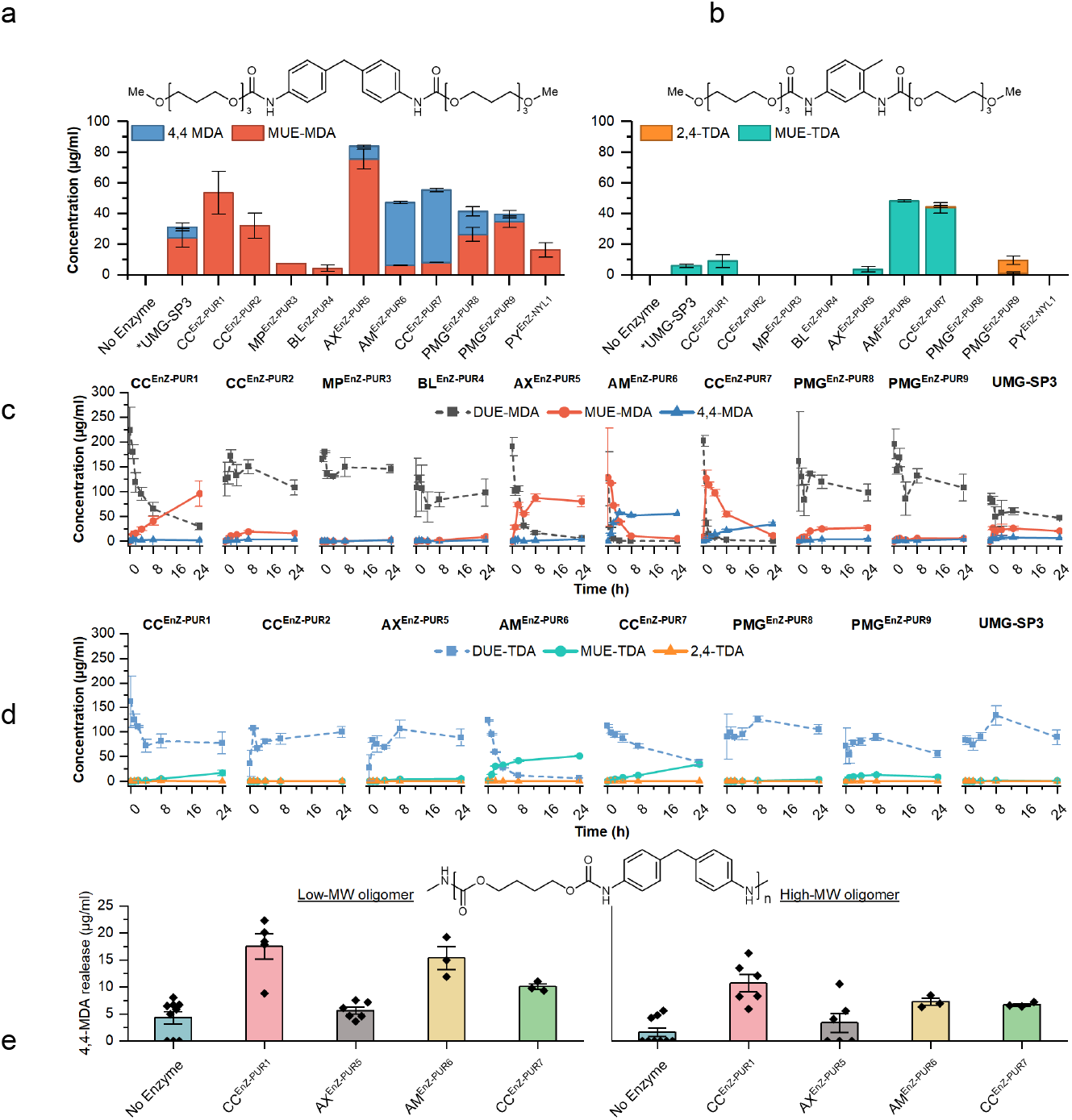
Activity of identified PURases on polyurethane. HPLC analysis of degradation products of DUE-MDA and DUE-TDA. Reactions with 0.1mg/ml substrates were stopped after 48h at 35℃ and quantified. ( a) Quantification of the two reaction products from either a single cleavage (MUE-MDA, red) or double cleavage (4,4-MDA, blue). Data is shown as mean of 3 replicates with error bars as SE. (b) Quantification of the two reaction products from either a single cleavage (MUE-TDA, teal) or double cleavage (2,4-TDA, orange). Data is shown as mean of 3 replicates with error bars as SE. Time resolved degradation of (c) DUE-MDA and (d) DUE-TDA. Reactions with 0.1 mg/ml DUE-TDA or DUE-MDA and 500 nM enzyme were sampled at 0, 1, 2, 4, 8, and 24h. Reactions were stopped by acetonitrile (1:1 dilution, 100 µg/mL theoretical HPLC concentration). Reactions were run according to the observed optimum activity on PUR-fluorophore. (e) MDA Oligomer degradation. Activity of enzymes on two different sizes of insoluble PUR oligomers as measured by the release of 4,4-MDA monomers on HPLC. 7 days, 1µM or 5 µM of enzyme, 35oC. Note: detection limit for 4,4MDA was 3.6 µg/ml. Results are shown as means of three replicates with SE unless otherwise shown.

**Figure 3.**
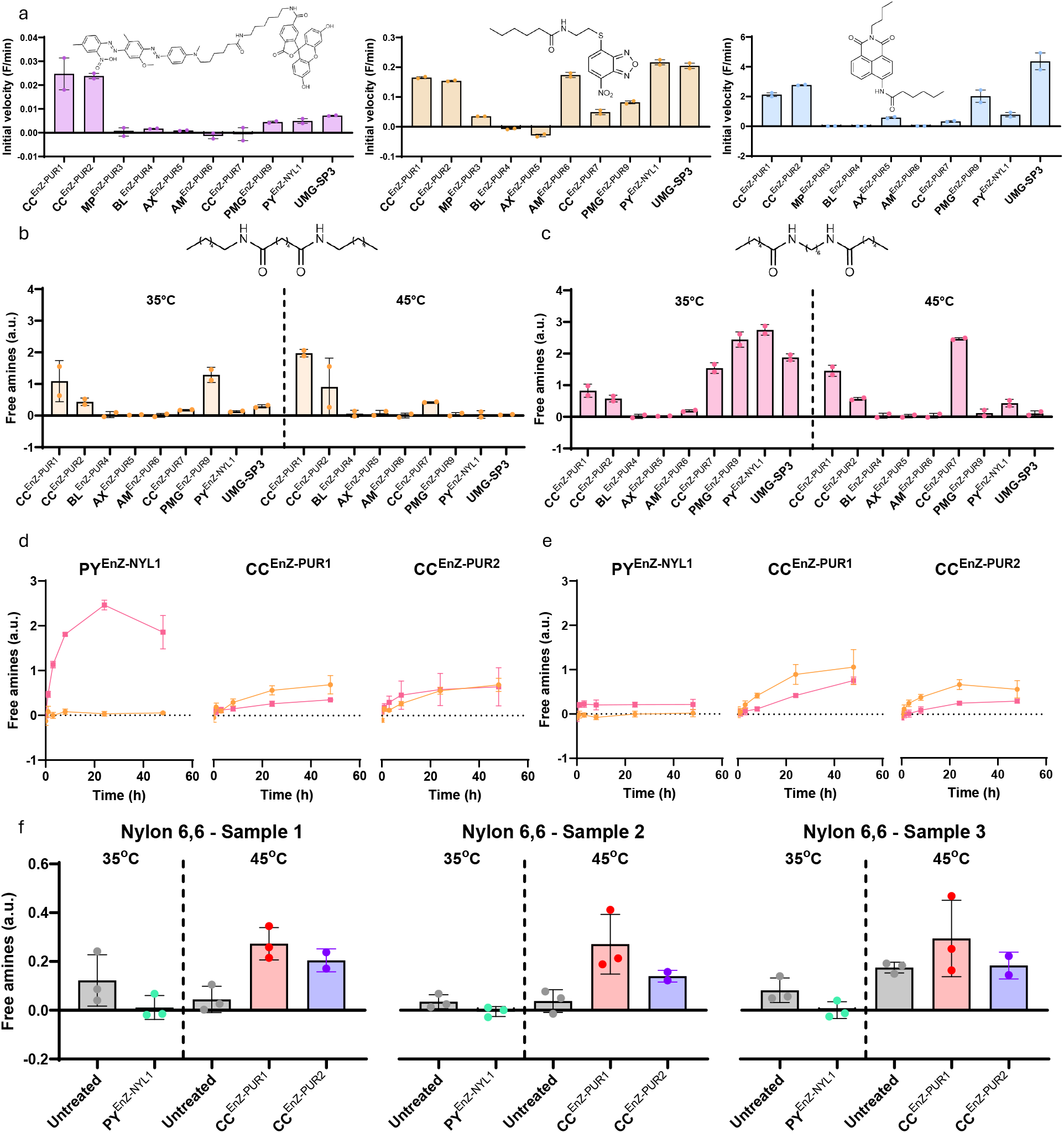
Activity of identified enzymes on nylon substrates. (a) Initial rates of probe-based screening of potential NYLONases. Substrate negative controls have been subtracted. Activity of selected enzymes on insoluble nylon dimers (b) CO-mimic and (c) N-mimic at 35°C and 45°C. Time-course analysis at (d) 35°C and (e) 45°C of the activity on insoluble nylon dimers CO-mimic (orange) and N-mimic (pink) of PY^EnZ-NYL1^, CC^EnZ-PUR1^ and CC^EnZ-PUR2^. (f) Release of free amine in real nylon 6,6 threads and textiles samples in presence or absence of PY^EnZ-NYL1^, CC^EnZ-PUR1^ and CC^EnZ-PUR2^ at their optimal temperatures. Sampel 1 (threads) and sample 3 (textile) present a fiber diameter of 20-30 μm, while sample 2 (textile) has a fiber diameter of 30-40. Negative controls of the buffer and the corresponding enzymes have been subtracted. Data is shown as mean and errors as standard error of the mean. Release of free amine groups is measured as increase of NBD-Cl absorbance at 474 nm.

Fewer of these enzymes degraded DUE-TDA, with much lower conversion rates compared to DUE-MDA (**Fig. 2b**). Five out of the ten enzymes tested showed release of up to ∼50 µg/ml MUE-TDA at 35℃ after 48h (**Fig. 2b**). Only two enzymes (CC^Enz-PUR7^ and PMG^EnZ-PUR9^) showed detectable release of the 2-4-TDA monomer (**Fig. 2b**). As observed with DUE-MDA, many of the enzymes only degrade to the intermediate MUE-TDA. PMG^EnZ-PUR9^ released the highest detectable amount of 2-4 TDA at 8.2 µg/ml. AM^Enz-PUR6^ and CC^Enz-PUR7^ released 48.2 ± 1.2 µg/ml and 43.7.5 ± 5.7 µg/ml MUE-TDA respectively, significantly more than the previously characterised PURase UMG-SP3 (6.0 ± 1.8 µg/ml) (**Fig. 2b**). Given the many different compositions of PUR in use, enzymes with selective activities towards different formulations such as TDI- and MDI-based PUR are needed. The different activities on DUE-TDA and DUE-MDA are thus useful starting points for evolution of selective PURases.

To evaluate the performance of each candidate PURase over 24h, we repeated the assay at each enzyme’s individual pH and temperature optima (**Fig. S6, Fig. 2cd**). AM^Enz-PUR6^ was the most productive at converting both DUE-MDA to 4,4 MDA and DUE-TDA to MUE-TDA, with 55.4 ± 1.7 and 51.3 ± 1.6 µg/mL detected over 24h. After 1h, DUE-MDA had been fully converted to MUE-MDA; the release of 4,4-MDA by AM^Enz-PUR6^ at 4h was significantly higher than for UMG-SP3 after 24h (**Fig. 2b**). However, AM^Enz-PUR6^ did not release any 2,4-TDA despite almost full conversion of DUE-TDA to MUE-TDA (**Fig. 2d**). CC^Enz-PUR7^ and AX^EnZ-PUR5^ also showed enhanced conversion of most DUE-MDA converted after just 4h (**Fig. 2c**). CC^Enz-PUR7^ released up to 34.3 ± 6.1 µg/ml 4,4-MDA while AX^EnZ-PUR5^ only produced 4.0 ± 1.0 µg/ml 4,4-MDA from MUE-MDA after 24h, despite significant formation of MUE-MDA (**Fig. 2c)**. Similar profiles with a fast initial conversion to MUE-MDA were seen in PMG^EnZ-PUR8^, CC^EnZ-PUR2^, and UMG-SP3, with conversion slowing down after 8h (**Fig. 2c**). Interestingly, although most of the enzymes capable of hydrolysing PURp1 also showed activity on DUE-MDA, their catalytic efficiencies differed markedly between the two substrates. Notably, AM^EnZ-PUR6^ was the most efficient on DUE-MDA, yet the least efficient on PURp1, highlighting that fluorophores are to be used only for initial screening.

Analogous to the DUEs, we designed two nylon 6,6 insoluble di-amide mimics with either the amine N,N’-(hexane-1,6-diyl)-dihexanamide or the carbonyl group (N^1^,N^6^-dihexyladipamide) in a central position within the molecule. These substrates dissolved in 100% DMSO at 60°C but were turbid in aqueous suspension. However, 7 days incubation of 2 mM of substrate with 1 μM of CC^EnZ-PUR1^, CC^EnZ-PUR2^, CC^EnZ-PUR7^, PMG^EnZ-PUR9^ or PY^EnZ-NYL1^ effectively degraded at least one of the nylon mimics (**Fig. 3bc**) at levels exceeding UMG-SP3, as demonstrated by the increase of free amines reported by NBD-Cl (**Fig. S8**) and a decrease of sample turbidity even at room temperature (**Fig. S9**). AX^EnZ-PUR5^ and BL^EnZ-PUR4^ showed no activity against these mimics (**Fig. 3bc**). All active enzymes showed activity levels (**Fig. 3bc**) that correlated with their thermostability (**Fig. S6**). PY^EnZ-NYL1^ and, to a lower extent, CC^EnZ-PUR7^ differentiated between the mimics, with a significant degradation of the N-mimic (**Fig. 3c**) but almost no effect on the CO-mimic (**Fig. 3b**).

Using NBD-Cl to monitor formation of free amine groups, we found PY^EnZ-NYL1^ to be more active than CC^EnZ-PUR1^ and CC^EnZ-PUR2^ (**Fig. 3d-e**), releasing levels of free amines within 3 h that CC^EnZ-PUR1^ and CC^EnZ-PUR2^ only achieved after 48 h (**Fig. 3d**). The high promiscuity of these latter proteins might lead to lower substrate affinity than PY^EnZ-NYL1^.

### Enzymatic degradation of insoluble oligomers and polymers

Plastic mimics such as our fluorescent probes and substituted carbamates such as DUE-TDA and DUE-MDA are only the first step in confirming plastic-degrading activity. Substrate design needs careful consideration. Commonly used substrates such as Impranil® DLN and other PUR polyesters are more likely to be degraded via the more labile ester bond. Therefore, we turned to hydrolysis of soluble oligomers that are chemically identical to the PUR and nylon as well as longer insoluble oligomers of PUR. Two different insoluble MDA based oligomers were synthesized and confirmed by Gel Permeation Chromatography and DOSY-NMR to be a low-MW oligomer and a high-MW oligomer (average molecular weight of ∼20 and ∼50 kDa, respectively), with distinct shift in retention times between the two oligomers (**Fig. S10** and **Fig. S11ab**). Trace amounts of short fragments >3 kDa was shown by MALDI-TOF mass spectrometry, but GPC confirms the majority is larger oligomers (**Fig. S10)**. After incubation of our best five PURases for 7 days at 35°C with the oligomers, all enzymes showed increased 4,4-MDA release on average compared to the control samples with 3-9 replicates tested (**Fig. 2e)**. CC^EnZ-PUR1^ showed the highest and most significant 4,4-MDA release on both the insoluble low-MW and high-MW oligomer, with 17.5 ± 2.3 µg/ml and 10.7 ± 1.6 µg/ml respectively, with 4.3 ± 1.1 µg/ml and 1.6 ± 0,8 µg/ml released for the same reactions in the absence of added enzyme. AM^EnZ-PUR6^ also released 15.4 ± 2.2 µg/ml 4,4-MDA monomer on the low-MW oligomer (**Fig. 2e)**. To our knowledge, this is the first observation of the degradation of insoluble high-MW oligomers of MDA-PUR. AX^EnZ-PUR5^ and CC^EnZ-PUR7^ exhibited minor qualitative release of 4,4′-MDA; however, activity did not consistently exceed background levels, potentially due to limited enzyme stability over the course of the assay and replicate values falling below the 3.6 µg/mL detection limit.

Finally, we tested PY^EnZ-NYL1^, CC^EnZ-PUR1^ and CC^EnZ-PUR1^ enzymes on three nylon textile samples composed of nylon 6,6 (**Fig. S11cde**). After incubation for 2 weeks at the enzymes’ optimal temperature, CC^EnZ-PUR1^ exhibited the most pronounced degradation of nylon textile samples, as reported by the increase of free amines in the solution (**Fig. 3f**). CC^EnZ-PUR2^ also increased the soluble free amine content in most of the tested samples except for one of the textile samples, where no clear differences could be observed compared to untreated samples (**Fig. 3f**). Despite its high efficiency on the nylon dimer, PY^EnZ-NYL1^ did not show any significant release of free amine into the solution (**Fig. 3f**). Nevertheless, the low NBD-Cl absorbance obtained from textile samples compared to the N- and CO-mimics, demonstrate the slow kinetics. The limited activity of PY^EnZ-NYL1^ is likely associated with its significantly lower stability, implying unfolding during the assay. Consistent with this, the most thermostable enzyme (CC^EnZ-PUR1^) is the most effective (**Fig. 3f** and **Fig. S6**).

## Conclusion

Using our FACS-based screening platform, we isolated 29 different bacterial species with either carbamate or amide cleavage, from which we identified eight enzymes by purification from bacterial lysates followed by MS. In addition, we discovered four enzymes using homology searches either within our discovered strains genomes or in published databases. Among our twelve isolated enzymes, ten were able to cleave DUE-MDA, five of them to the unsubstituted compound 4,4-MDA. Six were able to hydrolyse DUE-TDA, with only two releasing 2,4-TDA monomers, suggesting that TDA-based PUR represents a more recalcitrant substrate. Nine enzymes were active on at least one nylonase probe, five significantly degraded insoluble nylon mimics and CC^EnZ-PUR1^ and CC^EnZ-PUR2^ released free amines from nylon 6,6 textile samples. We also show monomer release from insoluble oligomers of MDA-based PUR. To our knowledge, this study is the first reported example of enzymes capable of hydrolysing longer chains of PUR and nylon and provides a platform for future optimization campaigns to develop enzymatic solutions to plastic degradation through *in silico* discovery and protein engineering.

## Acknowledgements

EnZync is generously supported by Challenge grant NNF22OC0072891 from the Novo Nordisk Foundation. We are deeply grateful to the rest of the EnZync consortium members for constructive and helpful feedback. Flow cytometry and cell sorting were performed at the FACS Core Facility, Aarhus University, Denmark. We are grateful to Bjarke Donslund for constructive discussions and Victoria Otzen for expert help in collecting samples. We thank Alexander Zelikin for GPC access.

## Author contributions

M.K.B., A.M and D.E.O. – Conceptualization; M.K.B., A.M., S.P.-D., R.G., N.C.P., B.N.I., M.C., M.B.J., A.S., A.R.P., I.K.C., C.R. and C.S. – Investigation; M.K.B., A.M., S.P.-D., R.G., Formal analysis; M.K.B., A.M., S.P.-D. and R.G. – Methodology; T.R.W., V.G., H.W., A.S and D.E.O. – Resources; A.S., V.G. and D.E.O – Supervision; A.S. and D.E.O. – Funding Acquisition; D.E.O. – Project Administration; M.K.B., A.M., S.P.-D., R.G., N.C.P., and D.E.O. Writing original draft; M.K.B., A.M., S.P.-D., R.G., N.C.P., M.B.J., A.S., A.R.P., A.S. and D.E.O. – Writing review.

