## Supplementary Information for "Environmental identification of novel enzymes against heteroatomic plastics"

### Environmental prospecting identifies novel enzymes targeting different heteroatomic plastics

#### Supplementary Information

### Methods

#### Synthesis of fluorescent probes and other substrates

##### PURase probe PURp1

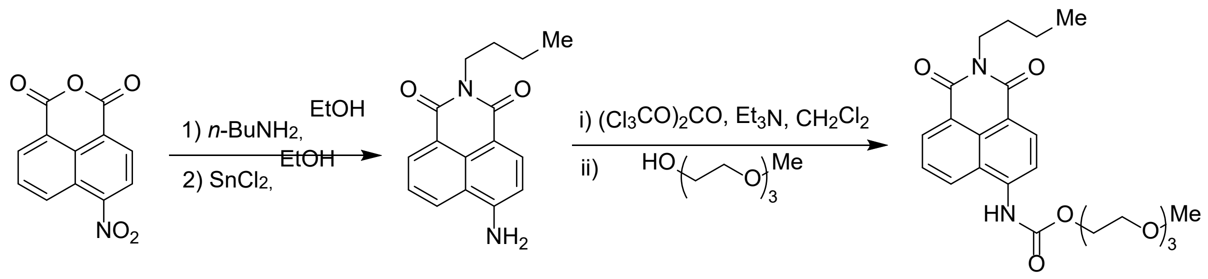


2-(2-(2-methoxyethoxy)ethoxy)ethyl (2-butyl-1,3-dioxo-2,3-dihydro-1H-benzo[de]isoquinolin-6-yl)carbamate (PURp1) was synthesized in three steps. Reaction mixtures, intermediate, and products were kept from light sources by protecting the glassware with aluminum foil and the lights in the fume hood were off when possible.

To make 2-butyl-6-nitro-1H-benzo[de]isoquinoline-1,3(2H)-dione**,** *n*-Butylamine (TCI Chemicals, 0.4 mL, 4.12 mmol, 1.03 equiv) was added dropwise to a solution of the 4-nitro-1,8-naphthalic anhydride (Apollo Scientific, 0.97 g, 4.00 mmol, 1.00 equiv) in ethanol (96 %, 50 mL). After 5 h reaction at 70 °C and stirring (200 rpm) [the substrate is fully solubilized after 1-2 h after which slight precipitation occurs], the reaction mixture was slowly cooled under continuous stirring (150 rpm). The precipitate was collected by filtration and washed with ethanol and further dried under vacuum to afford the title product as a light brown solid (357.6 mg). The filtrate was concentrated onto silica under reduced pressure and purified on a Büchi Pure C-850 FlashPrep (12 g column, heptane:EtOAc 100:0 to 50:50) to afford the title product (490.1 mg) as a pale yellow solid. The two solids were combined (71 %).

**$^1$H NMR (400 MHz, $CDCl_3$)**: δ 8.84 (dd, J = 8.7, 1.1 Hz, 1H), 8.77 – 8.66 (m, 2H), 8.41 (d, J = 7.9 Hz, 1H), 7.99 (dd, J = 8.8, 7.3 Hz, 1H), 4.24 – 4.15 (m, 2H), 1.79 – 1.67 (m, 2H), 1.46 (h, J = 7.4 Hz, 2H), 0.99 (t, J = 7.3 Hz, 3H).

**$^{13}$C NMR (101 MHz, $CDCl_3$)**: δ 163.5, 162.6, 132.6, 130.1, 129.9, 129.4, 129.3, 127.2, 124.1, 123.8, 123.2, 40.8, 30.2, 20.5, 14.0.

Next, to make 6-amino-2-butyl-1H-benzo[de]isoquinoline-1,3(2H)-dione (the cleaved PUR fluorophore), a solution of the 2-butyl-6-nitro-1H-benzo[de]isoquinoline-1,3(2H)-dione (1.70 g, 6.30

mmol, 1.00 equiv) in ethanol (96%, 25 mL) under nitrogen atmosphere was added $SnCl_2$ (3.60 g, 19.0 mmol, 3.00 equiv) at 0 °C under stirring (300 rpm). The reaction mixture was then heated to reflux and left to stir continuously under an atmosphere of nitrogen for 18 h. The mixture was cooled, and water was added slowly (25 mL). The mixture was quenched with $NaHCO_3$ (5 wt%, 50 mL), extracted with EtOAc (5 x 25 mL), and the combined organic phase was dried over $Na_2SO_4$, and concentrated under reduced pressure to afford the crude mixture.

The crude was concentrated onto celite under reduced pressure and purified on a Büchi Pure C-850 FlashPrep (12 g column, heptane:EtOAc 70:30 to 30:70 in a 25 min ramp) to afford the title product (0.8376 g, 50 %) as an orange solid.

**$^{1}$H NMR (400 MHz, DMSO-d6)** δ 8.59 (dd, J = 8.4, 1.2 Hz, 1H), 8.39 (dd, J = 7.3, 1.1 Hz, 1H), 8.17 (d, J = 8.4 Hz, 1H), 7.62 (dd, J = 8.4, 7.3 Hz, 1H), 7.42 (s, 2H), 6.82 (d, J = 8.4 Hz, 1H), 4.04 – 3.94 (m, 2H), 1.62 – 1.50 (m, 2H), 1.40 – 1.24 (m, 2H), 0.90 (t, J = 7.3 Hz, 3H).

**$^{13}$C NMR (101 MHz, DMSO-d6)**: δ 163.8, 162.9, 152.7, 133.9, 131.0, 129.7, 129.3, 124.0, 121.8, 119.4, 108.2, 107.6, 29.9, 19.9, 14.0.

Finally, to synthesise 2-(2-(2-methoxyethoxy)ethoxy)ethyl (2-butyl-1,3-dioxo-2,3-dihydro-1H-benzo[de]isoquinolin-6-yl)carbamate (PURp1), triethylamine (1.3 mL, 9.39 mmol, 3.00 equiv) was added dropwise to a solution of 6-amino-2-butyl-1H-benzo[de]isoquinoline-1,3(2H)-dione (0.84 g, 3.13 mmol, 1.00 equiv) in $CH_2Cl_2$ (30 mL) in a 100 mL round-bottomed flask charged with oval stir bar (400 rpm) under a nitrogen atmosphere. Next, a solution of trisphosgene (0.46 g, 0.157 mmol, 0.50 equiv) in $CH_2Cl_2$ (10 mL) [HCl fumes were formed] was added dropwise. The mixture was allowed to stir continuously for 2 h before triethylene glycol monomethyl ether (1.5 mL, 9.39 mmol, 3.00 equiv) was added at room temperature. The mixture was allowed to stir continuously at room temperature for 18 h.

The reaction mixture was quenched by slow addition of $NaHCO_3$ (5 wt%, 20 mL), transferred to a separation funnel, washed with additional $NaHCO_3$ (5 wt%, 3 x 25 mL), dried over $Na_2SO_4$, filtered, concentrated onto celite under reduced pressure, and purified on a Büchi Pure C-850 FlashPrep (12 g column, heptane:EtOAc 70:30 to 0:100 in a 25 min ramp + 5 min with EtOAc) to afford the title product (1.20g, 83 %) as a yellow solid.

**$^{1}$H NMR (400 MHz, $CDCl_3$)**: δ 8.60 (dd, J = 7.3, 1.0 Hz, 1H), 8.54 (d, J = 8.3 Hz, 1H), 8.32 (d, J = 8.3 Hz, 1H), 8.25 (dd, J = 8.6, 1.1 Hz, 1H), 7.90 (s, 1H), 7.73 (dd, J = 8.5, 7.3 Hz, 1H), 4.46 – 4.39

(m, 2H), 4.20 – 4.11 (m, 2H), 3.84 – 3.77 (m, 2H), 3.77 – 3.61 (m, 5H), 3.59 – 3.52 (m, 2H), 3.34 (s, 3H), 1.76 – 1.64 (m, 2H), 1.52 – 1.37 (m, 2H), 0.97 (t, J = 7.4 Hz, 3H).

**$^{13}$C NMR (101 MHz, $CDCl_3$)**: δ 164.3, 163.8, 153.4, 139.3, 132.6, 131.3, 129.0, 126.6, 126.4, 123.5, 123.1, 117.9, 116.7, 72.1, 70.8, 70.7 (2C), 69.3, 65.1, 59.1, 40.3, 30.3, 20.5, 14.0.

**HRMS (ESI+)**: m/z calcd. for $C_{24}H_{31}N_2O_7$ $[M+H]^+$: 459.2126, found 459.2121.

#### DUE-MDA and MUE-MDA

These were synthesized as described in Rotilio *et al.*[1].

#### DUE-TDA

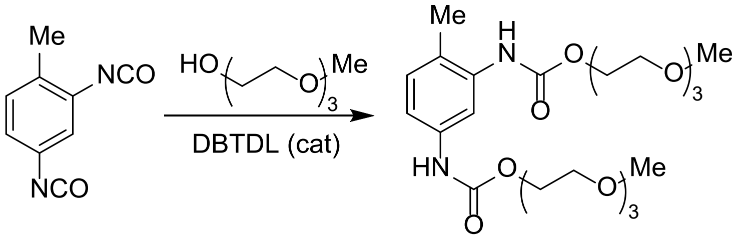


Under a nitrogen atmosphere, tolylene-2,4-diisocyanate (1.43 mL, 10 mmol, 1.00 equiv) was added to a mixture of triethylene glycol monomethyl ether (16 mL, 100 mmol, 10.0 equiv) and dibutyl tin dilaurate (DBTDL) (32 mg, 0.05 mmol, 0.01 equiv) in a 50 mL round-bottomed flask charged with oval stir bar (500 rpm). The mixture was stirred for 24 h. The reaction mixture was transferred to a separation funnel, diluted with EtOAc (50 mL) and washed with water (5 x 50 mL, mixed with appr. 1 V/V% brine), HCl (0.1 M, 3 x 50 mL), brine (50 mL). The organic phase was then dried above $Na_2SO_4$ and filtered before being concentrated under reduced pressure to afford a crude yellow oil. The mixture was purified on a Büchi Pure C-850 FlashPrep (12 g column, (EtOAc:$Et_3N$ 99:1):MeOH 100:0 to 96:4 in a 12 min ramp) to afford the title product (3.2 g, 64%) as a pale yellow oil.

**$^{1}$H NMR (400 MHz, $CDCl_3$)**: δ 7.77 (s, 1H), 7.20 (d, J = 7.7 Hz, 1H), 7.09 – 6.94 (m, 2H), 6.57 (s, 1H), 4.34 – 4.25 (m, 4H), 3.82 – 3.59 (m, 16H), 3.57 – 3.50 (m, 4H), 3.38 – 3.32 (m, 6H), 2.17 (s, 3H).

**$^{13}$C NMR (101 MHz, $CDCl_3$)**: δ 153.43 (2C), 136.72, 136.21, 130.75, 122.04 114.26, 111.07 71.94, 71.92, 70.60 (4C), 64.35, 64.13, 59.05, 17.08.

**HRMS (ESI+)**: m/z calcd. for $C_{23}H_{39}N_2O_{10}$ $[M+H]^+$: 503.2599, found 503.2599.

#### MUE-TDA

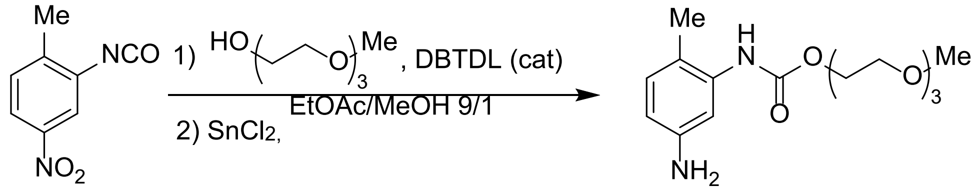


2-(2-(2-methoxyethoxy)ethoxy)ethyl (5-amino-2-methylphenyl)carbamate (MUE-TDA) was synthesized in two steps. Under a nitrogen atmosphere, 2-methyl-5-nitrophenyl isocyanate (1.00 g, 5.61 mmol, 1.00 equiv) was added to a mixture of triethylene glycol monomethyl (9.0 mL, 56 mmol, 10.0 equiv) and DBTDL (one drop) in a 40 mL vial charged with oval stir bar (600 rpm). The mixture was heated to 60 °C and then stirred continuously for 2 days. The reaction mixture was transferred to a separation funnel, diluted with EtOAc (200 mL) and washed with water (5 x 100 mL), HCl (0.1 M, 1 x 150 mL), brine (50 mL). The organic phase was then dried above $Na_2SO_4$ and filtrated before concentration under reduced pressure to afford a crude yellow oil (1.7 g). The crude mixture was purified on a Büchi Pure C-850 FlashPrep (12 g column, 5 min ramp heptane:EtOAc 70:30 to 55:45 in a 5 min ramp, then 5 min at heptane:EtOAc 55:45, then heptane:EtOAc 55:45 to 80:20 in a 10 min ramp) to afford a yellow oil (1.12 g, 58 %).

**$^1$H NMR (400 MHz, $CDCl_3$)**: δ 8.64 (s, 1H), 7.76 – 7.68 (m, 1H), 7.20 (d, *J* = 8.3 Hz, 1H), 7.11 (d, *J* = 5.0 Hz, 1H), 4.27 (tt, *J* = 4.4, 1.8 Hz, 2H), 3.71 – 3.64 (m, 2H), 3.65 – 3.53 (m, 6H), 3.50 – 3.43 (m, 2H), 3.26 (s, 3H), 2.27 (s, 3H).

**$^{13}$C NMR (101 MHz, $CDCl_3$)**: δ 153.4, 146.7, 137.0, 134.7, 130.7, 118.2, 115.2, 71.7, 70.4, 70.4, 70.3, 69.1, 64.6, 58.8, 17.9.

Then, under a nitrogen atmosphere, tin(II)chloride (3.88 g, 20.4 mmol, 14.0 equiv) was added to a solution of 2-(2-(2-methoxyethoxy)ethoxy)ethyl (2-methyl-5-nitrophenyl)carbamate (0.50 g, 1.45 mmol, 1 equiv) in EtOAc/MeOH 9/1 V/V (20 mL) in 40 mL vial under stirring (600 rpm) at room temperature and allowed to stir continuously for 5 days. The reaction mixture was quenched using a sodium carbonate solution (10 mL, concentrated). The precipitating white tin compounds were removed by filtration and washed with EtOAc (3 x 50 mL). The combined organic phases were washed with brine (2 x 10 mL), dried over $Na_2SO_4$ and concentrated under reduced pressure to obtain the crude product. The crude mixture was purified on a Büchi Pure C-850 FlashPrep (12 g column, heptane:EtOAc 20:80 to 0:100 in a 6 min ramp and then 7 min with pure EtOAc) to afford 2-(2-(2-methoxyethoxy)ethoxy)ethyl (5-amino-2-methylphenyl)carbamate as a colorless oil (407 mg, 89%).

**$^{1}$H NMR (400 MHz, CDCl$_3$)**: δ 7.29 (s, 1H), 6.89 (d, $J$ = 8.0 Hz, 1H), 6.58 (s, 1H), 6.36 (dd, $J$ = 8.0, 2.4 Hz, 1H), 4.34 – 4.26 (m, 2H), 3.83 – 3.45 (m, 12H), 3.36 (s, 3H), 2.11 (s, 3H).

**$^{13}$C NMR (101 MHz, CDCl$_3$)**: δ 153.6, 145.2, 136.6, 131.0, 116.8, 110.9, 107.5, 72.0, 70.7, 70.6 (2C), 69.6, 64.3, 59.1, 16.9.

**HRMS (ESI+)**: m/z calcd. for $C_{15}H_{25}N_2O_5$ $[M+H]^+$: 313.1758, found 313.1766.

#### *N,N'*-dihexyladipamide (CO-mimic)

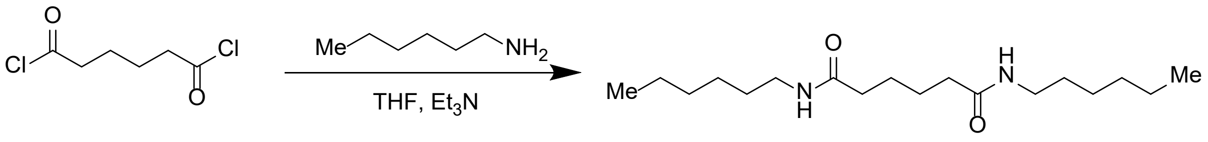


Protocol inspired by Zhou *et al.*[2]:

In a 250 mL round bottom flask, hexylamine (5.3 mL, 40 mmol, 2.00 equiv) and triethylamine (6.7 mL, 48 mmol, 2.40 equiv) were dissolved in THF (100 mL). The solution was cooled to 0 °C and adipoyl chloride (2.9 mL, 20 mmol, 1.00 equiv) was added dropwise, the system was warmed up to room temperature and stirred continuously for 5 days. The solvent was removed under reduced pressure. To this residue was added water (50 mL), the slurry was filtered, and solid was washed by THF, and dried under vacuum to obtain a white solid (4.00 g, 64 %).

**$^{1}$H NMR (400 MHz, CDCl$_3$)** δ 5.69 (t, J = 5.8 Hz, 2H), 3.23 (q, J = 6.6 Hz, 4H), 2.15 (t, J = 7.6 Hz, 4H), 1.61 (p, J = 7.5 Hz, 4H), 1.48 (p, J = 6.7 Hz, 4H), 1.39 – 1.21 (m, 12H), 0.88 (t, J = 6.8 Hz, 6H).

**$^{13}$C NMR (101 MHz, CDCl$_3$)** δ 173.4, 39.0, 37.0, 31.6, 29.6, 26.1, 25.7, 22.5, 14.1.

The NMR spectra are in accordance with literature data[3].

#### *N,N'*-(hexane-1,6-diyl)dihexanamide (N-mimic)

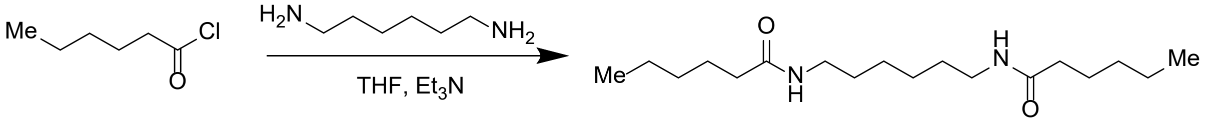


Protocol inspired by[2]:

In a 250 mL round bottom flask, 1,6-hexanediamine (2.6 mL, 20 mmol, 1.00 equiv) and triethylamine (6.7 mL, 48 mmol, 2.40 equiv) were dissolved in THF (100 mL). The solution was cooled to 0 °C and hexanoyl chloride (5.6 mL, 40 mmol,21.00 equiv) was added dropwise, the system was warmed up to room temperature and stirred continuously for 5 days. The solvent was removed under reduced pressure. To this residue was added water (50 mL), the slurry was filtered, and solid was washed by

THF, and dried under vacuum to obtain a white solid (4.55 g, 73 %). **$^{1}$H NMR (400 MHz, $CDCl_3$)** δ 5.70 (t, *J* = 6.0 Hz, 2H), 3.29 – 3.17 (m, 4H), 2.22 – 2.09 (m, 4H), 1.61 (p, *J* = 7.4 Hz, 4H), 1.55 – 1.42 (m, 4H), 1.39 – 1.21 (m, 12H), 0.88 (t, *J* = 6.8 Hz, 6H). **$^{13}$C NMR (101 MHz, $CDCl_3$)** δ 173.4, 39.0, 37.0, 31.6, 29.6, 26.1, 25.7, 22.5, 14.1.

The NMR spectra are in accordance with literature data[2]:

#### Nylonase Probe 1 (NYLp1)

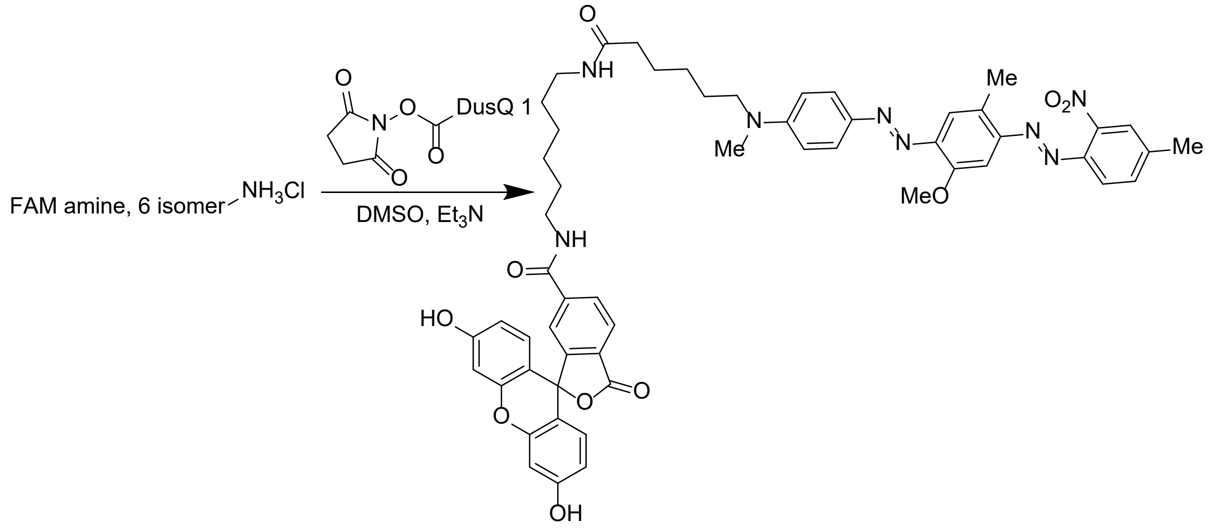


The DusQ 1 NHS ester (Lumiprobe, 25.0 mg, 39.7μmol, 1.00 equiv) was solubilized in 400 μL DMSO in a vial and added slowly to a mixture of FAM amine, 6-isomer (Lumipobe, 20.3 mg, 39.7 μmol, 1.00 equiv), triethylamine (27.7 μL, 199 μmol, 5.00 equiv), and 400 μL DMSO in a vial at room temperature under stirring (400 rpm). The mixture was purged with nitrogen for 1 min before the vial was closed, wrapped with aluminium foil, and the reaction mixture was allowed to stir at room temperature for 5 days. The mixture was diluted with EtOAc (50 mL), transferred to a separation funnel, and washed with a $NH_4Cl$ solution (5 wt%, 5 x 50 mL). The organic phase was dried over $Na_2SO_4$, filtered, and concentrated under reduced pressure to afford a purple solid. This crude mixture was diluted in $CH_2Cl_2$, concentrated onto celite, and purified on a Büchi Pure C-850 FlashPrep (4 g column, EtOAc in a 16 min ramp) to afford the title product (24.5 mg, 67 %) as a purple solid. **$^{1}$H NMR (400 MHz, Acetone)** δ 9.25 (s, 2H), 8.23 (dd, *J* = 8.0, 1.6 Hz, 1H), 8.18 – 8.10 (m, 1H), 8.02 (d, *J* = 8.0 Hz, 1H), 7.87 – 7.72 (m, 5H), 7.64 (t, *J* = 6.7 Hz, 1H), 7.55 (d, *J* = 3.2 Hz, 1H), 7.37 (d, *J* = 3.0 Hz, 1H), 7.22 – 7.11 (m, 1H), 6.84 – 6.73 (m, 4H), 6.71 – 6.57 (m, 4H), 3.95 (s, 3H), 3.50 – 3.40 (m, 2H), 3.35 – 3.26 (m, 2H), 3.12 (q, *J* = 6.5 Hz, 2H), 3.08 – 3.03 (m, 3H), 2.67 (s, 3H), 2.54 (s, 3H), 2.15 (t, *J* = 7.3 Hz, 2H), 1.69 – 1.56 (m, 4H), 1.53 – 1.44 (m, 2H), 1.45 – 1.21 (m, 8H).

Impurities present below 1.4 ppm and near 3.6 ppm. Water assumed to be peak at 3.00-2.85. **$^{13}C$ NMR (101 MHz, Acetone)** δ 173.6 – 173.3 (m), 168.9, 165.9 – 165.8 (m), 160.5, 155.9, 154.1, 153.3, 152.9, 151.5, 148.5, 146.26, 145.0, 143.7, 143.4, 142.2, 134.4, 133.5, 130.2, 130.1, 129.7, 126.5, 125.5, 125.0, 123.4, 120.2, 119.4, 113.5, 112.2, 111.1, 103.4, 100.28, 56.52, 52.8, 40.4 – 40.2 (m), 39.5 – 39.3 (m), 38.8, 36.6 – 36.5 (m), 27.4, 27.2, 27.0, 26.8, 26.3, 21.1, 16.8. **HRMS (ESI+)**: m/z calcd. for $C_{55}H_{57}N_8O_{10}$ $[M+H]^+$: 989.4192, found 989.4206.

#### Nylonase Probe 2 (NYLp2)

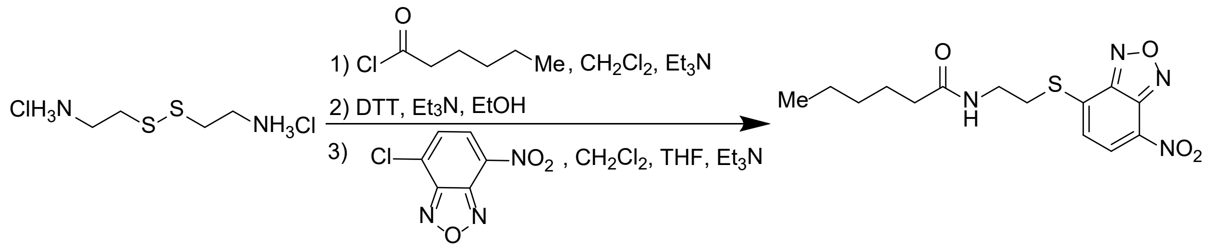


N-(2-((7-nitrobenzo[c][1,2,5]oxadiazol-4-yl)thio)ethyl)hexanamide (NYLp2) was synthesized in 3 steps.

First a dry 100 mL flask was charged with a large stir bar, cystaminium dichloride (2.25 g, 10 mmol, 1.0 equiv) and $CH_2Cl_2$ (30 mL) was added followed by triethylamine (6.97 mL, 50 mmol, 5.0 equiv). Subsequently, a solution of hexanoyl chloride (3.49 mL, 25 mmol, 2.5 equiv) in $CH_2Cl_2$ (15 mL) was added dropwise over 30 min with precipitation of the triethylammonium hydrochloride salt. The solution was stirred for 1 h at room temperature and then quenched with water (30 mL). The organic phases were washed with 0.1 M HCl (3 x 50 mL), water (50 mL), 5% $NaHCO_3$ (3 x 50 mL), water (50 ml) and brine (50 ml) and finally dried over $MgSO_4$, filtered and concentrated under reduced pressure to give a white solid.

The amide product was approximately 90% pure by $^1H$ NMR and used without further purification.

Then the disulfide (1.00 g, 2.87 mmol, 90% purity, 1.0 equiv) was reduced with dithiothreitol (0.796 g, 5.17 mmol, 1.8 equiv) dissolved in degassed absolute ethanol (20 mL) and degassed triethylamine (0.02 mL, 0.144 mmol, 0.05 equiv) was added. The solution was stirred overnight at room temperature and then diluted with ethyl acetate (50 mL) and water (50 mL). The organic phase was washed with 0.1 M HCl (2 x 50 mL), water (3 x 50 mL) and brine (50 mL), dried over $MgSO_4$, filtered, and concentrated under reduced pressure to give a transparent oil, which was used without further purification.

The thiol product was approximately 80% pure by $^{1}H$ NMR, with the major impurity being the disulfide starting material, and was used without further purification.

Finaly, 4-Chloro-7-nitrobezofurazan (0.100 g, 0.5 mmol, 1.0 equiv) was dissolved in degassed dichloromethane/tetrahydrofuran (1:1, 2 mL), and the thiol (0.088 g, 0.5 mmol, 80% pure, 1.0 equiv) was added. Triethylamine (0.084 mL, 0.6 mmol, 1.2 eq) was added and the mixture was stirred at room temperature for 1 hr. The crude mixture was then poured into diethyl ether (50 mL), washed with 5% $NaHCO_3$ (2 x 50 ml), water (2 x 50 mL) and brine (50 mL). The organic phase was dried over $MgSO_4$, filtered and evaporated onto celite, which was subjected to dry column chromatography (Pentane/EtOAc, 10:1 to 1:10 with 10% increments in EtOAc and 50 mL per fraction). The desired compound was obtained as an orange solid (30 mg, 17% yield). $R_f$(product, EtOAc) = 0.7, $R_f$(product, 1:1 EtOAc/pentane) = 0.2.

**$^{1}$H NMR (80 MHz, $CDCl_3$)** δ 8.42 (d, $J$ = 7.92 Hz, 1H), 7.69 (d, $J$ = 8 Hz, 1H), 6.25-6.15 (m, 1H), 3.70-3.30 (m, 4H), 2.23 (t, $J$ = 7.62 Hz, 2H), 1.80-1.00 (m, 12H), 0.87 (t, $J$ = 5.9 Hz, 4H).

**$^{13}$C NMR (20 MHz, $CDCl_3$)** δ 174.3, 142.5, 140.3, 132.9, 131.3, 125.5, 121.6, 38.4, 36.6, 31.5, 30.5, 25.4, 22.5, 14.0

#### N-(2-butyl-1,3-dioxo-2,3-dihydro-1H-benzo[de]isoquinolin-6-yl)hexanamide (Nylonase probe 3, NYLp3)

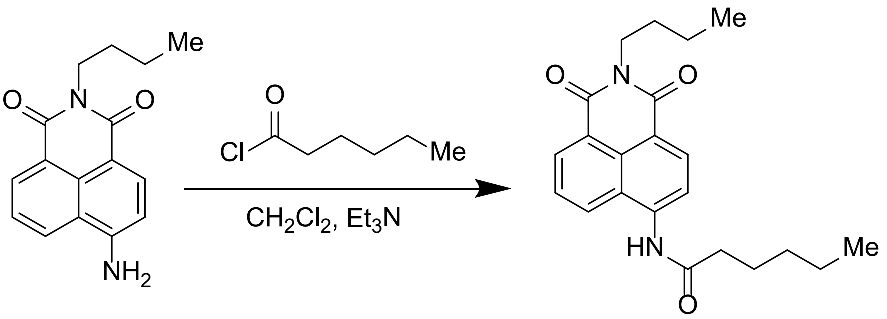


In a 20 mL amber vial was 6-amino-2-butyl-1H-benzo[*de*]isoquinoline-1,3(2*H*)-dione (110 mg, 0.40 mmol, 1.0 equiv) and trimethylamine (0.07 mL, 0.52 mmol, 1.3 equiv) were dissolved in $CH_2Cl_2$ (5 mL). Hexanoyl chloride (0.06 mL, 0.43 mmol, 1.05 equiv)) in $CH_2Cl_2$ (5 mL) was added dropwise, the system was stirred continuously for 7 days. Crude mixture was concentrated onto celite under reduced pressure and purified on a Büchi Pure C-850 FlashPrep (25 g column, heptane:EtOAc 90:10 for 5 min, 90:10 to 75:25 in a 10 min ramp, then 75:25 for 9 min, then 75:25 to 0:100 in a 2 min ramp, and then pure EtOAc for 3 min) to afford the title product as a beige solid (31.3 mg, 21 %).

**$^{1}$H NMR (400 MHz, DMSO)** δ 10.34 (s, 1H), 8.67 (dd, *J* = 8.6, 1.1 Hz, 1H), 8.51 (dd, *J* = 7.2, 1.0 Hz, 1H), 8.46 (d, *J* = 8.2 Hz, 1H), 8.29 (d, *J* = 8.2 Hz, 1H), 7.88 (dd, *J* = 8.5, 7.2 Hz, 1H), 4.03 (t, *J* = 7.3 Hz, 2H), 2.57 (t, *J* = 7.5 Hz, 2H), 1.74 – 1.55 (m, 4H), 1.41 – 1.31 (m, 6H), 0.96 – 0.87 (m, 6H).

**$^{13}$C NMR (101 MHz, DMSO)** δ 172.6, 163.5, 163.0, 140.3, 131.7, 130.9, 129.3, 128.3, 126.4, 124.1, 122.3, 117.5, 36.2, 31.0, 29.7, 24.8, 22.0, 19.9, 13.8.

**HRMS (ESI+)**: m/z calcd. for $C_{22}H_{27}N_2O_3$ $[M+H]^+$: 367.2017, found 367.2013.

#### Oligomers of MDI and butanediol

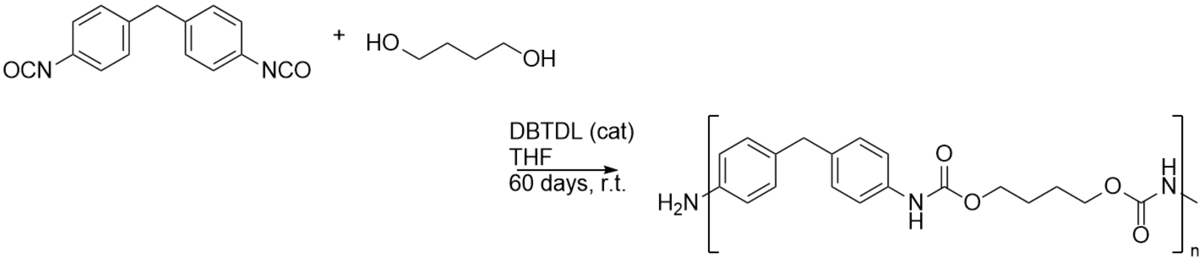


Linear polymeric PUR was made from from 4,4’-methylene diphenylisocyanate (MDI) and 1,4-butanediol (BDO) in various sizes.

To a 50 mL flame dried round bottomed flask equipped with a magnetic stir bar was added either 2.500 gram (high-MW oligomer) or 1.650 gram (Low-MW oligomer) of a fresh unopened 4,4’-methylene diphenylisocyanate, followed by 25 mL dry THF stored over molecular sieves. 0.885 µL volume of 1,4-butanediol is measured using a micropipette, and added in one step, followed by 10 µL dibutyl tin dilaurate. The reaction solution was sealed using a glass stopper and stirred for 60 days at room temperature. The reaction was opened to air and left to dry out over a day. Each sample was then dissolved in between 100 mL DMSO at 50 °C overnight. The dissolved sample was precipitated in a 1 L conical flask with demineralized water, stirred at 500 rpm using a large rod magnetic bar making a deep vortex in the water. 1 drop of polymeric solution was added into the vortex and evaluated. If necessary, the solution was diluted with additional DMSO until the drop of polymeric solution into vortex resulted in polymer being dispersed in the water as medium long threads. The dispersed polymer was isolated *via* vacuum filtration, redissolved once more in DMSO, and precipitated once more. Upon final isolation, the polymer was dried at 80 °C under vacuum for 24 hours. Resulting in white powder in near quantitative yield.

**$^{1}$H NMR (80 MHz, DMSO-*d6*)** δ 9.52 (bs, 2H), 7.33 (d, 4H), 7.15 (d, 4H), 4.12 (bs, 4H), 3.80 (bs, 2H), 1.72 (bs, 4H).

Additional peaks observed for the Low-MW oligomer amine end-terminal samples:

**$^{1}$H NMR (80 MHz, DMSO-*d6*)** δ 8.52 (bs), 6.75 (d), 6.50 (d).

#### 1H-NMR End group Analysis:

End-group analysis was made for the Low-MW oligomer using the integral from 6.80 to 7.65 ppm (corresponding to 8 protons) as backbone and the integral from 6.6 to 6.3 ppm as terminal group (corresponding to 4 protons). The resulting end-group analysis corresponds to an average DP of 33.4 and an average molecular weight of 11.436 g/mol.

End group analysis was not possible on the high-MW oligomer.

#### SEC-MALS/GPC

The dried samples (Low-mw oligomer and High-mw oligomer) (~3 mg) were dissolved in DMF (1 mL) and filtered through a 0.2 µM pore size filter. Size exclusion chromatography was performed on a system consisting of an Agilent 1260 Infinity II Isocratic HPLC Pump, an Agilent 1260 Infinity II VWD UV-Vis detector, a Wyatt miniDAWN 3-angle static light scattering detector and a Shodex RI 501 refractive index detector, using DMF with 10 mM lithium bromide as mobile phase. The setup was equipped with a three-column system consisting of PLgel columns (Agilent Technologies, particle size 5 µm, length 300 mm, inner diameter 7.5 mm) with pore sizes of 100000, 500 and 50 Å, respectively, resulting in effective molecular ranges spanning from 10-450, 5-250 and >1.5 kDa. Measurements were conducted at T = 40 °C, with a flowrate of 1 mL/min. The system was calibrated with a 200 kDa PS standard. Average molecular weights and polydispersity were calculated assuming 100% mass recovery, applying a d*n*/d*c* of 0.105[4].

Low-MW oligomer: $M_n$= 19766 (±5.29%) Da, $M_w$= 25169 (±4.09%) Da, PDI = 1.27 (±6.69%)

high-MW oligomer: $M_n$= 52282 (±2.59%) Da, $M_w$= 81574 (±1.22%) Da, PDI = 1.56 (±2.86%)

#### DOSY-NMR

DOSY NMR was performed using a Magritek Spinsolve 80 Multi-X ULTRA spectrometer via a PGSTE pulse sequence with carbon decoupling (PGSTE CDEC) and the Spinsolve software for analysis. The samples <5 mg were dissolved in deuterated DMSO.

From the DOSY experiment analysis diffusion constants where determined, and from these an approximate molecular weight was determined using the molecular weight calculation software provided by the university of Warwick[5].

Low-MW oligomer: Diffusion constant = 2.322E-11 $m^2s^{-1}$ corresponding to 21,818 Da in DMSO.

High-MW oligomer: Diffusion constant = 1.338E-11 $m^2s^{-1}$ corresponding to 54,950 Da in DMSO.

#### MALDI-TOF

Matrix assisted laser desorption/absorption ionization time-of-flight (MALDI-TOF) mass spectrometry experiments were performed using a Bruker Daltonics Autoflex Speed spectrometer, an AnchorChip target plate, and trans-2-[3-(4-tert-Butylphenyl)-2-methyl-2- propenylidene]malononitrile (DCTB) in MeCN as matrix (1ug/mL). Samples were prepared by dissolving 1 ug material in 1 mL DMF. Following addition of 1 uL sample onto the plate, DMF was evaporated by heating the plate to 80 C for 1 min. After allowing the plate to cool, 1 uL matrix solution was added. N

#### Isolation of polyurethanese- and nylonases-active bacteria

Bacteria with plastic-degrading enzymes were isolated from diverse environments. Polyurethane samples were obtained from an indoor tropical zoo (Randers Regnskov, Denmark) while both polyurethane and nylon were sampled from equatorial landfills (Mwakirunge Garbage Dump, Kenya). Polymer identity was verified by attenuated total reflectance Fourier-transform infrared spectroscopy (4500 Series Portable FTIR Spectrometer, Agilent, USA). Environmental enrichments were prepared by incubating industrial garden compost (Lisbjerg Recycling Station) with polyurethane foam, elastane or nylon for 3-5 months. *Galleria mellonella* (wax worm) larvae were fed polyurethane foam for 7 days after which the guts were extracted. The microorganisms were extracted by vortexing for 30 min at 2,700 rpm in PBS with 0.5 % Tween 20. The microbial suspensions were sequentially centrifuged to pellet and remove larger particles (1,000 g, 10 min) and to pellet the microorganisms (10,000 g, 10 min). The pellets were resuspended in PBS and filtered through a 70 µm mesh (Flowmi Cell Strainer, SP Bel-Art, USA).

Microorganisms were screened for their hydrolytic activity using Fluorescence-Assisted Cell Sorting (Bigfoot Cell Sorter, Invitrogen, USA) at 37°C with 100 µM fluorescent probes (PURp1 and NYLp1). For PURases screening, uncleaved and cleaved probes were detected by excitation/emission at 349/473 nm and 445/525 nm, respectively. To avoid bacterial aggregates, microorganisms with a higher fluorescence signal of uncleaved probe were excluded from sorting. For nylonase screening, the released fluorophore was detected at 445/525 nm. The cell population with the highest fluorescent signal of cleaved or released probe was sorted onto an LB agar plate and grown for 7 days at 40°C (**Fig. S1d** and **S2c**).

To determine whether the enzyme was secreted or cell-associated, the isolates were cultured to high optical density ($OD_{600}$) at 40ºC. The supernatant was sterile filtered while the cells were resuspended in PBS. The fluorescence of the cleaved probe PURp1 (450/550 nm) was measured in a FLUOstar Omega Microplate Reader (BMG Labtech, Ortenberg, Germany) over 24 h at 40°C for both cells and supernatant. To compare relative activities between the isolates, the enzyme activity was normalized to $OD_{600}$.

Bacterial genomes were extracted using the DNeasy PowerSoil Pro kit (Qiagen, Hilden, Germany) and sequenced (Novogene, Beijing, Chine). Raw reads were trimmed using Trimmomatic and assembled with Spades[6] v. v. 4.0.0 or 4.1.0 and quality controlled using QUAST[7] v. 5.2.0 or 5.3.0 and CheckM2[8] v. 1.0.2 or 1.1.0 and assembled with Prokka[9] v. 1.14.6. Species were identified using ANI with GTDB-tk[10] to the nearest species or genus and plotted on a phylogenetic tree to display the taxonomic diversity. Due to IP concerns and ongoing enzyme discovery efforts, isolates with unidentified enzymes are indicated on the class level. Full species names will be made available upon future enzyme identification.

#### Enzyme Identification from Native Lysates by chromatography

Isolated bacterial strains were grown in LB to $OD_{600} \approx 0.6\text{-}0.8$ and lysed with 0.5mg/ml DNase and 5mg/ml lysozyme either by sonication or bead beating in PowerBoad Pro Tubes (Qiagen). Enzymes of interest were isolated from the supernatant by rounds of chromatography (Anion Exchange (AEX), Size exclusion (SEC), and hydrophobic interaction (HIC)) on an ÄKTA pure™ chromatography system (Cytiva, Marlborough, USA). Activity of eluted fractions were determined with 25µM PURase or nylonase probe (ex/em. 450/550 nm) and measured on a Varioskan™ LUX multimode microplate reader (Thermo Scientific). AEX was performed using a 5mL HiTrap Q HP (Cytiva) using a linear gradient of 100% sample buffer (50 mM Tris-HCl, pH 8) to 100% elution buffer (50 mM Tris-HCl,

1M NaCl, pH 8) over 5-10 column volumes at 3-5ml $min^{-1}$. SEC was performed on a Superdex™ 200 Increase 10/300 GL column (Cytiva) using a flow rate of 0.5 ml $min^{-1}$ using 50mM sodium phosphate, pH 7 running buffer. HIC was performed using either a HiTrap 1ml butyl HP or a HiPrep Octyl FF 16/10 (Cytiva) column according to manufacturer recommendations at 1ml $min^{-1}$ with a linear gradient of 100% binding buffer (50 mM $NaPO_4$, 1.5M $(NH_4)_2SO_4$, pH 7) to 100% elution buffer (50 mM $NaPO_4$, pH 7) over 5-10 column volumes.

After two to three rounds of column chromatography, the enzymes were purified further using zymography and PURp1. The active fractions were loaded onto native Mini-Protean TGX gels (Bio-Rad, Hercules, USA), run in native buffer (25 mM Tris-HCl, 192 mM glycine, pH 8) and stained with 50 µM PURp1 for 15-20 min. The active enzyme band was identified in a Typhoon Biomolecular Imager (Cytiva) by imaging the fluorescence of the cleaved probe (488/575 nm). The proteins were fixed (5 % acetic acid, 5 % ethanol) and the active bands were prepared for LC-MS/MS analysis using in-gel digestion. The in-gel digestion was performed as previously described with the minor modification of using 10 mM DTT and 30 mM iodoacetamide during the reduction and alkylation step[11]. Prior to the LC-MS/MS analysis the tryptic peptides were purified by solid phase extraction using Empore™ SPE Disks of C18 octadecyl packed in 10 µl pipette tips[12].

LC-MS/MS analyses were performed using an Easy nLC 1200 connected to a Tribrid Eclipse mass spectrometer (Thermo Fisher Scientific, Waltham, MA, USA). The samples were trapped on a pre-column (ReproSil-Pur C18-AQ 3 µm, Dr. Maisch GmbH, Germany). The peptides were eluted and separated on a 15 cm analytical column (75 µm i.d.) packed with ReproSil-Pur C18-AQ 3 µm in a pulled emitter. Peptides were eluted at a flow rate of 250 nL $min^{-1}$ using a 20 min gradient from 5% to 40% of solution B (0.1% formic acid and 90% acetonitrile). The collected Raw files were peak picked using RawConverter v1.2 and searched against the host proteome using an in-house Mascot 2.8.2 search engine (matrix science). Search parameters allowed one missed trypsin cleavage site with peptide tolerance and MS/MS tolerance set to 10 ppm and 50 mmu, respectively.

From the list of proteins identified in the digest, 2-4 enzymes from each isolate were selected for heterologous expression based on their annotated function. Enzymes were expressed in *Escherichia coli* BL21(DE3) using IPTG at 37°C and screened for their ability to cleave PURp1, NYLp1 or NYLp2. Enzymes with low expression levels ($AM^{Enz\text{-}PUR6}$ and $CC^{Enz\text{-}PUR7}$) were expressed in autoinduction media at 25°C.

#### Determination of optima

##### Melting point determination

0.5 mg/ml of each enzyme in 50mM phosphate pH 8, 50mM NaCl were heated from 25°C to 95°C at 1°C min$^{-1}$ in two replicates of two technical repeats on a Prometheus Panta nanoDSF (NanoTemper Technologies GmbH, Munich, Germany). The 350nm/330nm fluorescence ratio or turbidity were fitted to either a two- or three state equilibrium unfolding model in Moltenprot[42]. Data is shown as average of four repeats.

##### Determination of pH optimum

The optimal pH for each enzyme was recorded in the following buffers containing 100mM NaCl and 100mM sodium citrate (pH 4, pH 4.5, pH 5, pH 5.5, pH 6), BisTris propane (pH 6.5, pH 7, pH 7.5, pH 8, pH 8.5, pH 9, pH 9.5), CAPS ( pH 10, pH 10.5, pH 11), 30nM (CC$^{EnZ-PUR1}$, CC$^{EnZ-PUR2}$, BL$^{EnZ-PUR3}$, AX$^{EnZ-PUR5}$, PMG$^{EnZ-PUR8}$, PMG$^{EnZ-PUR9}$ PY$^{EnZ-NYL1}$ ) or 1µM (MP$^{EnZ-PUR4}$, AM$^{EnZ-PUR6}$, CC$^{EnZ-PUR7}$) of each enzyme was incubated for 15 min in assay buffer in a 96 well plate before 50µM PURp1 was added and activity was measured (ex./em. 450/550 nm, 12nm excitation band width, bottom optics, 37°C) with a Varioskan™ LUX multimode microplate reader (Thermo Scientific).

##### Determination of temperature optima

Enzymes were preincubated for 15 min at assay temperature before adding 25µM PURp1 probe. Fluorescence (ex./em 450/550 nm) was measured on a Cary Eclipse Fluorescence Spectrometer (Agilent, Santa Clara CA, United States) in 100mM BisTris-propane pH 9 with enzyme (30nM – 10µM depending on activity). Activity was determined as a slope of fluorescence intensity over the initial part of reaction. Results are reported as average of three replicates with substrate in assay buffer subtracted as background and normalized within each enzyme dataset.

##### Enzymatic degradation of DUE-MDA and DUE-TDA

Endpoint measurements: 500 nM of enzyme was incubated with 0.1mg/ml di-urethane ethylene toluenediamine (DUE-TDA) or di-urethane ethylene methylenedianiline (DUE-MDA) for 48h at 35°C in 100mM phosphate buffer pH 8 with 100mM NaCl.

Time courses: 500 nM of enzyme was incubated with 0.1mg/ml di-urethane ethylene toluenediamine (DUE-TDA) or di-urethane ethylene methylenedianiline (DUE-MDA) for 24h at the optimal temperature and pH according to screening on BIQTEG-Carbamate. Aliquots were taken at 0, 1, 2, 4, 8, and

24. The reaction was stopped by adding 100% AcN in a ratio 1:1. The aliquots were filtered through a 0.22µM PTFE filter and analysed on HPLC as above.

Standard curves of DUE-MDA (7.1ml), MUE-MDA (6.8ml), 4,4-MDA (6.2ml), DUE-TDA (6.1ml), MUE-TDA (5.5ml) and 2,4-TDA (4.3ml) were used to assign peaks and convert areas to concentrations. All standard curves could be described linearly with a $R^2 \geq 0.95$.

#### Enzymatic degradation of PUR oligomers and polymers

Assays were performed at 35°C in 100mM phosphate buffer pH 8 with 100mM NaCl. Enzyme concentrations were 1 µM for CC$^{\mathrm{EnZ\text{-}PUR1}}$, 1 µM for AX$^{\mathrm{EnZ\text{-}PUR5}}$, 5 µM for CC$^{\mathit{Enz}\mathrm{\text{-}PUR7}}$, 1 µM for AM$^{\mathit{Enz}\mathrm{\text{-}PUR6}}$, ~3-5 mg substrate per reaction with a final volume of 300µL. Before the reaction the oligomers were washed in 100% acetonitrile and dried completely under vacuum to ensure full removal of potential small soluble species.

#### HPLC analysis

All reactions were stopped by adding 100% acetonitrile (AcN) at a 1:1 ratio. Aliquots were filtered through a 0.22µM PTFE filter and analysed on HPLC on a Shimadzhu HPLC system using UV-absorbance at 240nm. 10µl of sample were injected onto a Zorbax Eclipse Plus C18 column 5µm particle size at 40°C and a flow rate of 1ml min$^{-1}$. The mobile phase A consisted of 5% acetonitrile (AcN) in miliQ and B of 95% Acetonitrile in milliQ water. The sequence consisted of a 2 min at 100% (A) followed by a gradient of 0-100% (B) for 8 min followed 1 min at 100% (B) and then a 30s gradient back to 100% (A). Finally, 3 min at 100% (A) was used to reequilibrate the column.

#### Screening for Nylonase potential

Nylonase activity was tested with three different nylon-mimics probes: N-(2-((E)-(4-((E)-(4-((6-((6-(3',6'-dihydroxy-3-oxo-3H-spiro[isobenzofuran-1,9'-xanthene]-6-carboxamido)hexyl)amino)-6-oxohexyl)(methyl)amino)phenyl)diazenyl)-5-methoxy-2-methylphenyl)diazenyl)-5-methylphenyl)-N-oxohydroxylammonium (NYL1), N-(2-((7-nitrobenzo[c][1,2,5]oxadiazol-4-yl)thio)ethyl)hexanamide (NYL2) and N-(2-butyl-1,3-dioxo-2,3-dihydro-1H-benzo[de]isoquinolin-6-yl)hexanamide (NYL3) (**Fig. S1**). The reaction was performed with 1 µM enzyme in a 96-well plate in 100 mM of bis Tris-propane pH 9.0 at 37°C under quiescent conditions. Probes were dissolved in 100% DMSO and added to a final concentration of 50 µM NYL1, 600 µM NYL2 or 50 µM NYL3 (final DMSO of 2.5, 6% and 2.5%, respectively). Fluorescence was measured in a Varioskan LUX Multimode Microplate Reader (Thermo Fisher Scientific, MA, USA) with ex/em 450/590 nm, 500/560 nm and 450/550 nm for NYL1, NYL2 and NYL3, respectively. Wells containing only the fluorophore were used as a negative control whose fluorescence was subtracted from treated samples. Initial rates were calculated as the slope of the florescence emission during the first 15 min of the reactions.

#### Activity on nylon dimers

As more realistic nylon-mimics we used a monoamine-based substrate CO-mimic $N^1$,$N^6$-dihexyladipamide, and a diamine-based substrate N-mimic, N,N'-(hexane-1,6-diyl)dihexanamide. Both designs contained two amide bonds that could be cleaved. 25 mM (7,81 mg/mL) of both dimers were resuspended in 100% DMSO and heated at 60°C until complete resuspension was achieved. Substrate was maintained at 60°C to remain soluble and only added to the reaction in the final step.

Nylon dimer degradation was performed in a 1,5 mL tube with 750 µL of 100 mM of bis Tris-propane pH 9.0 containing 1 µM of enzyme and 2 mM (0,64 mg/mL) of the dimer (added last, leading to an immediate increase of turbidity). Samples were then incubated in an Eppendorf PCMT ThermoMixer (Grant Instruments, Royston, UK) at either 35°C or 45°C and 500 rpm, after which the reaction was stopped by adding 750 µL of 100% acetonitrile and the sample stored at -20°C until measurement. Negative controls of the buffer and the different enzymes were used as reference.

For the screening assay, the samples were incubated for 7 days and at least two independent assays were performed. Due to substrate insolubility, during kinetic studies independent aliquots were used for each time point (0h, 1h, 3h, 8h, 24, and 48h). These assays were limited to the most active enzymes: $CC^{EnZ\text{-}PUR1}$ and $CC^{EnZ\text{-}PUR2}$ (selected for their capacity to degrade a variety of nylon mimics and improved activity at high temperatures) and $PY^{EnZ\text{-}NYL1}$ (selected for its specificity). $PMG^{EnZ\text{-}PUR9}$ was discarded at this point since its temperature resistance and activity did not exceed those the ones observed in the other candidates. Similarly, $CC^{EnZ\text{-}PUR7}$ was also discarded despite its higher stability compared to $PY^{EnZ\text{-}NYL1}$ due to its lower degrading capacity. Three independent tests were performed per conditions.

#### Degradation of nylon textile samples

The nylon degradation capacity of $CC^{EnZ\text{-}PUR1}$ and $CC^{EnZ\text{-}PUR2}$ and $PY^{EnZ\text{-}NYL1}$ was tested using 10-15 mg of textile samples composed by nylon 6,6 differing in the material stock and form: threads (sample 1) with 20-30 µm fibre diameter (**Fig. S11c**), and textiles (samples 2 and 3) with 30-40 µm (**Fig. S11d**) or 20-30 µm fibre diameter (**Fig. S11e**). To increase the surface exposure of the sample, the plastic was manually disaggregated prior to the assay. The reaction was performed in a solution of 500-750 µL of 100 mM of bis Tris-propane pH 9.0 at the optimal temperature of the enzymes, 45°C

($CC^{EnZ-PUR1}$ and $CC^{EnZ-PUR2}$) or 35°C ($PY^{EnZ-NYL1}$). The plastic was incubated in an Eppendorf PCMT ThermoMixer (Grant Instruments, Royston, UK) in presence or absence of 5 µM of the enzymes (0,25-0,3 mg/mL) for 2 weeks at 500 rpm. Once completed, the soluble fraction was recovered, and the reaction was stopped by adding of 100% acetonitrile in a 1:1 volume ration. Finally, the sample stored at -20°C until measurement.

#### Detection of free amines

Release of free amine was quantified with 4-Chloro-7-nitrobenzofurazan (NBD-Cl), previously re-suspended in 100% DMSO. Amine functionalisation was performed in a 96-well plate containing 100 µL of the sample, 5 mM of NBD-Cl and 50% DMSO, to a final volume of 200 µL. Changes on the absorbance spectrum was recorded for 1h by measuring from 400 nm to 600 nm in a CLARIOstar platereader (BMG LABTECH, Ortenberg, Germany). Optimal measurement differences were observed after 15 min at 474 nm. Intrinsic absorbance of buffer, enzymes and untreated dimer were subtracted.

### Supplementary Table S1

| Name | Application | Structure | Fluorescence |
|---|---|---|---|
| **PURp1** | Fluorescent PURase probe | 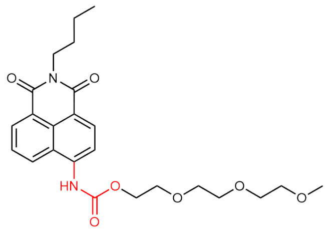 | Ex.: 450 nm<br>Em.: 550 nm |
| **NYLp1** | Fluorescent nylonase probe | 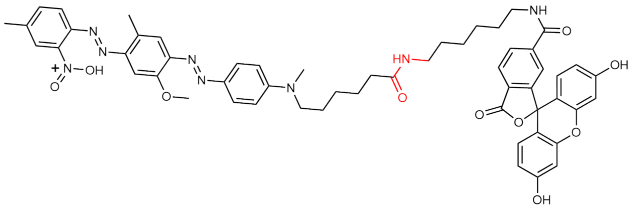 | Ex.: 450 nm<br>Em.: 590 nm |
| **NYLp2** | Fluorescent nylonase probe | 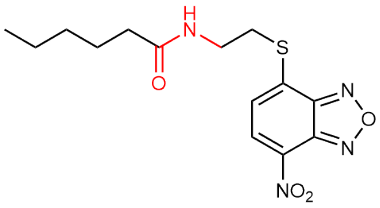 | Ex.: 500 nm<br>Em.: 560 nm |
| **NYLp3** | Fluorescent nylonase probe | 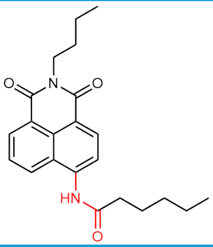 | Ex.: 450 nm<br>Em.: 550 nm |
| **DUE-MDA** | MDA-based soluble PUR mimic | 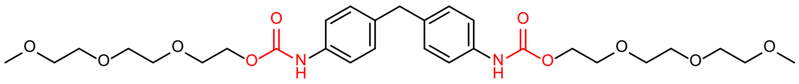 | N/A |
| **DUE TDA** | TDA-based soluble PUR mimic | 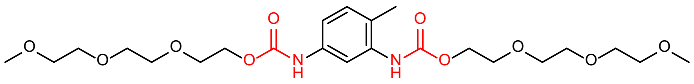 | N/A |
| **CO-mimic** | Nylon 6,6 dimer mimic | 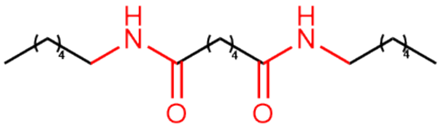 | N/A |
| **N-mimic** | Nylon 6,6 dimer mimic | 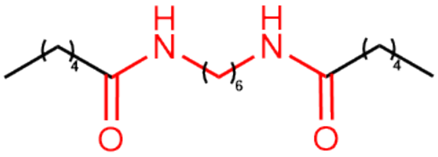 | N/A |
| **Low-MW PUR oligomer/ High-MW PUR oligomer** | Oligomeric forms of PUR | 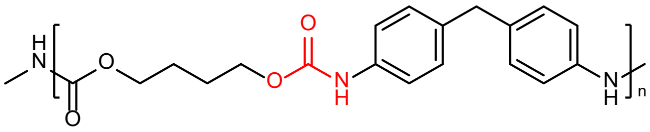 | N/A |

### Supplementary Table S2

| Enzyme | Purified from | Column #1 | Column #2 | Column #3 | Zymography |
|---|---|---|---|---|---|
| CC$^{EnZ\text{-}PUR1}$ | Lysate (soluble fraction) | AIX | SEC | HIC | NO |
| CC$^{EnZ\text{-}PUR2}$ | Lysate (soluble fraction) | AIX | SEC | - | YES |
| MP$^{EnZ\text{-}PUR3}$ | Homology to AX$^{EnZ\text{-}PUR5}$ | | | | |
| BL$^{EnZ\text{-}PUR4}$ | Secretome | SEC | AIX | HIC | NO |
| AX$^{EnZ\text{-}PUR5}$ | Lysate (soluble fraction) | AIX | SEC | HIC/AIX | YES |
| AM$^{Enz\text{-}PUR6}$ | Lysate (Soluble Fraction) | AIX | SEC | HIC | NO |
| CC$^{Enz\text{-}PUR7}$ | Lysate (Soluble Fraction) | AIX | SEC | HIC | NO |
| PMG$^{EnZ\text{-}PUR8}$ | Homology to CC$^{EnZ\text{-}PUR1}$ | - | - | - | - |
| PMG$^{EnZ\text{-}PUR9}$ | Homology to CC$^{EnZ\text{-}PUR1}$ | - | - | - | - |
| PY$^{EnZ\text{-}NYL1}$ | Homology to CC$^{EnZ\text{-}PUR1}$ | - | - | - | - |
| Hydrolase from *M. sciuri* | Lysate (soluble fraction) | AIX | SEC | HIC/AIX | YES |
| Amidase from *Metapseudomonas* sp. | Lysate (soluble fraction) | AIX | SEC | HIC | NO |

### Figure S1

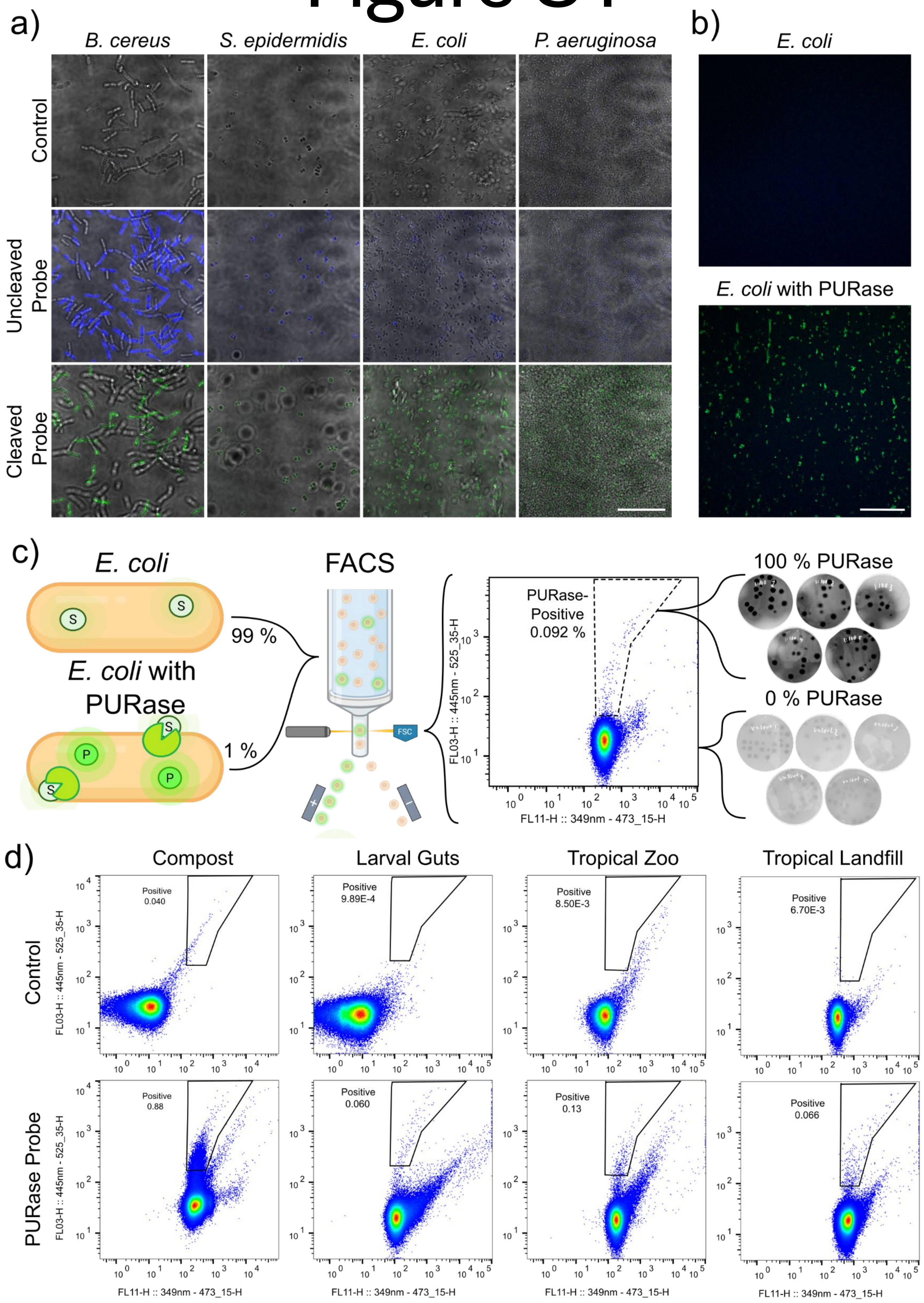


**Supplementary Figure S1. Verification of PURase probe for sorting by FACS**

a). Penetration of cleaved and uncleaved PURase probe (PURp1) into Gram-positive (*Bacillus cereus* and *Staphylococcus epidermidis*) and Gram-negative (*E. coli* and *Pseudomonas aeruginosa*) bacteria. Penetration of probe was verified by CLSM. Scalebar is 20 µm.

b) Staining of PURase-positive *E. coli* expressing recombinant PURase (UMG_SP2) with PURp1 probe. Scalebar is 20 µm.

c) FACS sorting of *E. coli* where 1 % of the population expresses recombinant PURase (UMG-SP2). The subpopulation with highest fluorescence of cleaved PURp1 was sorted onto agar plates with PURp1. The fluorescence of the cleaved PURp1 was imaged, confirming that all colonies on the sorted plates were PURase-positive (n=81). When the whole population was sorted onto plates, none of the colonies were PURase-positive (n=93).

d) Examples of FACS scatterplots from the four different PURase sources, with and without PURp1 probe. The signal of uncleaved and cleaved probe were plotted on each axis at 349nm – 473_15 and 445nm – 525_35, respectively. Gates indicate subpopulations with high signal of cleaved probe.

### Figure S2

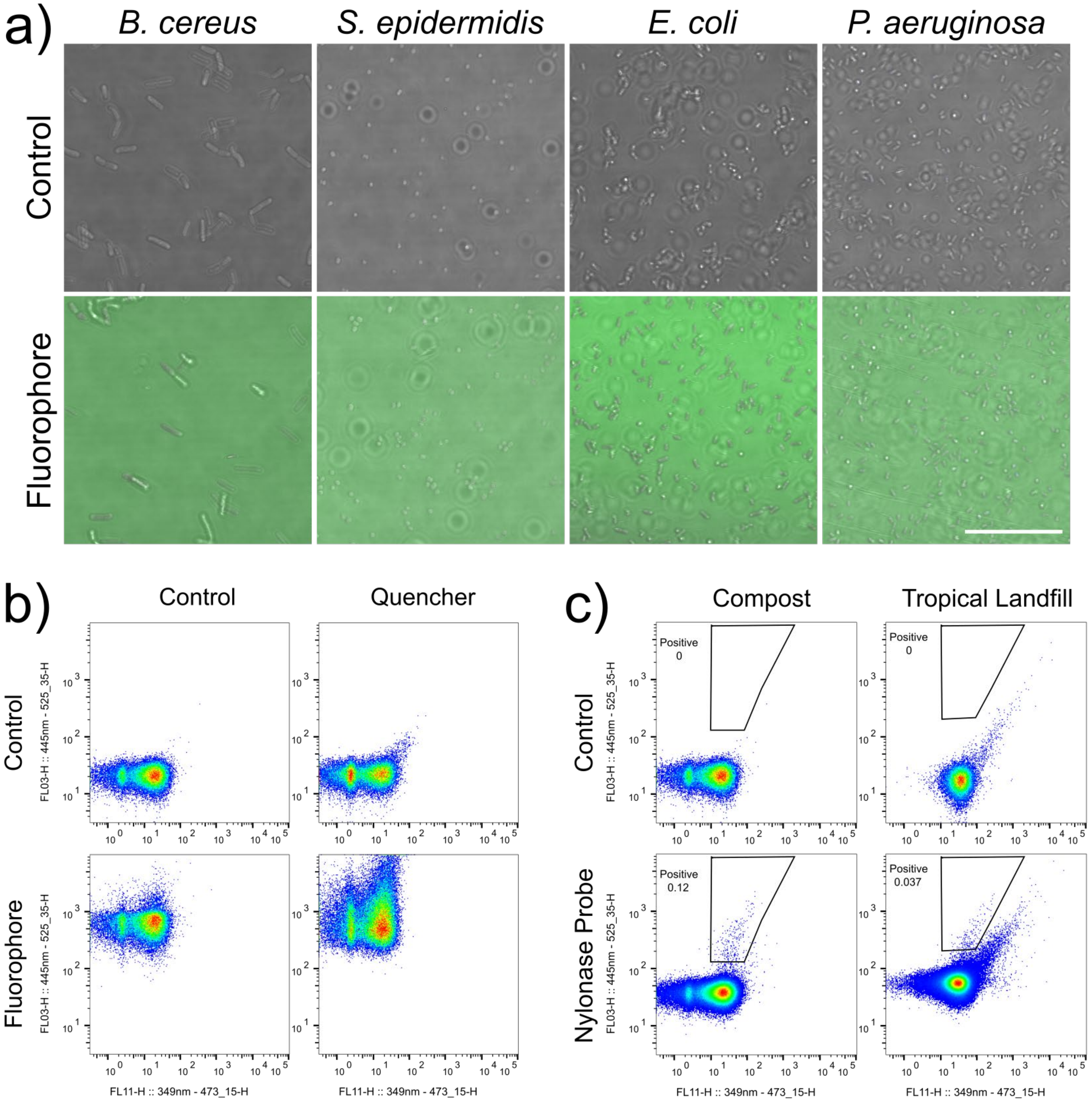


**Supplementary Figure 2. Verification of nylonase probe NYLp1 for sorting by FACS**
a). Lack of penetration by fluorophore released from cleaved nylonase probe (NYLp1) incubated with Gram-positive (*B. cereus* and *S. epidermidis*) and Gram-negative (*E. coli* and *P. aeruginosa*) bacteria. Lack of penetration was verified by CLSM. Scalebar is 20 µm.
b) FACS scatterplots of compost microbiome stained by NYLp1 fluorophore, quencher and both. The signal of released fluorophore (445nm – 525_35) is plotted along the y axis.
c) Examples of FACS scatterplots from the two different nylonase sources, with and without NYLp1. Gates indicate subpopulations with high signal of released fluorophore.

### Figure S3

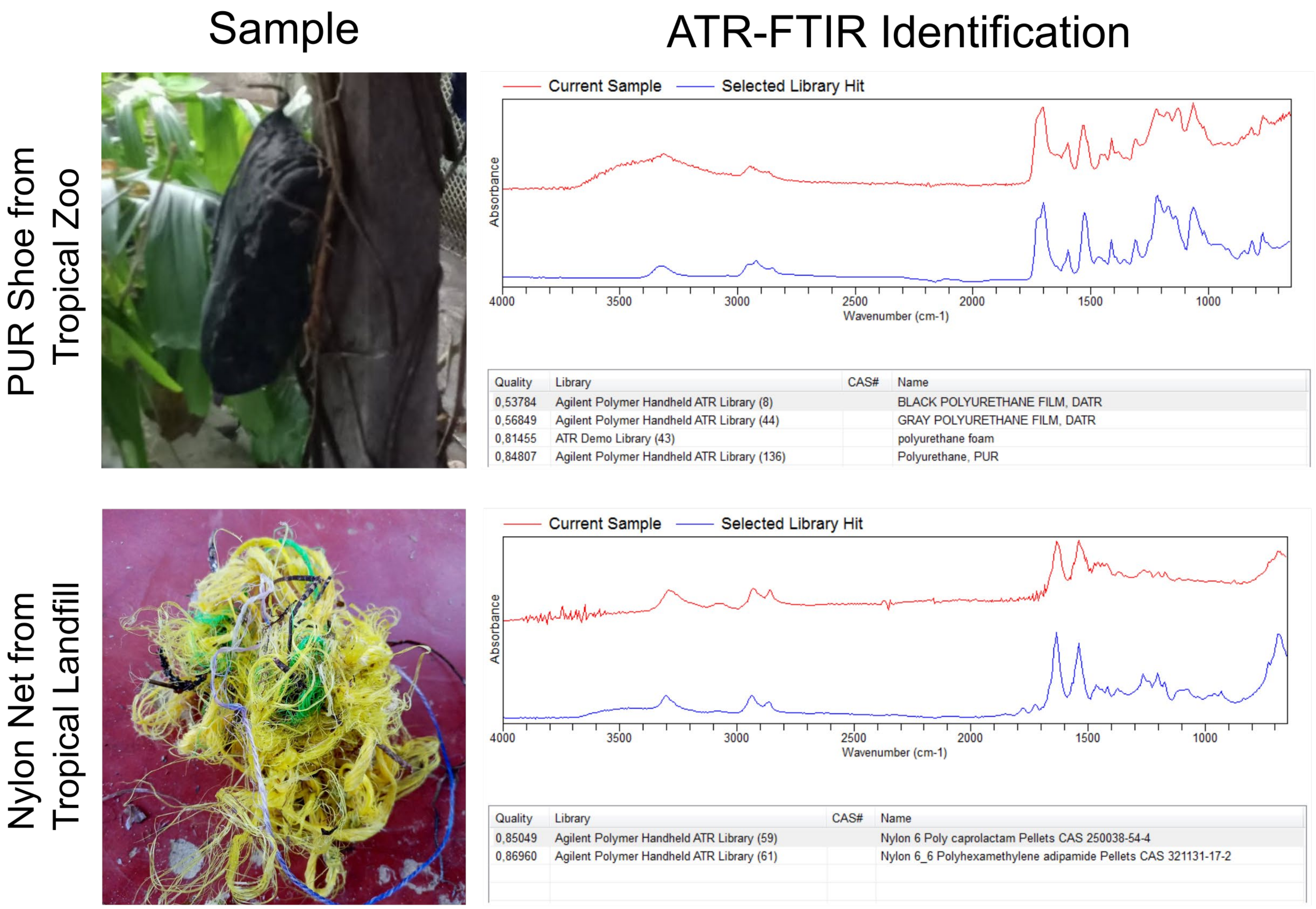


**Supplementary Figure 3. Identification of polymer composition of plastic samples by ATR-FTIR.**
Examples of plastic samples from the tropical zoo and landfill. The samples were confirmed to be PUR or nylon by ATR-FTIR and cross-referencing the spectra with a library.

### Figure S4

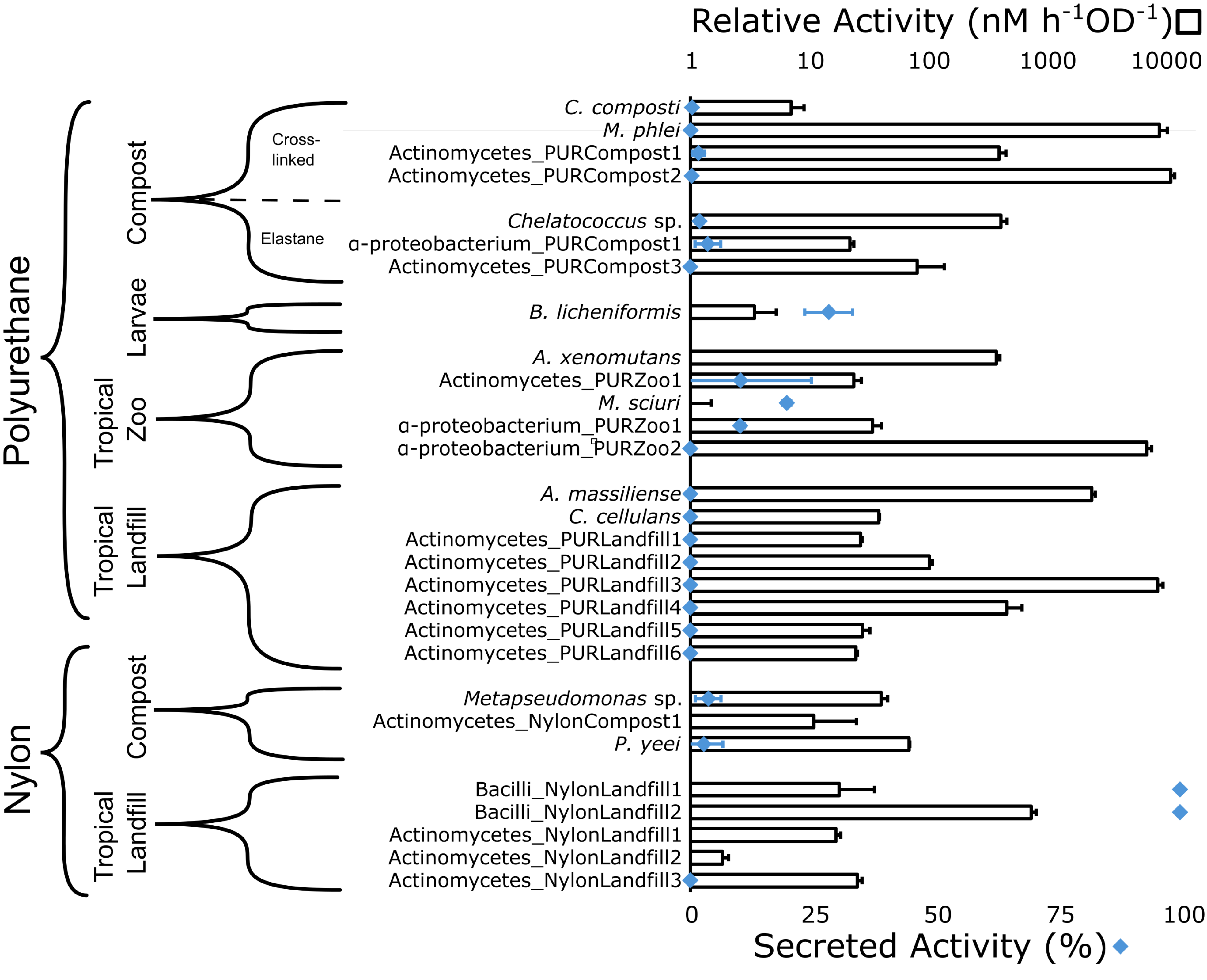


**Supplementary Figure 4. Overview of plasticase activity by environmental isolates.** Relative PURase- and nylonase activity of environmental isolates, adjusted for optical density (n=3). Activity is against PURp1 and NYLp1 probes, respectively. Brackets on the left indicate source of the isolate. Diamonds indicate the fraction of enzyme activity that was secreted to the supernatant (n=3). Bars with no diamonds showed slightly negative secreted activity.

### Figure S5

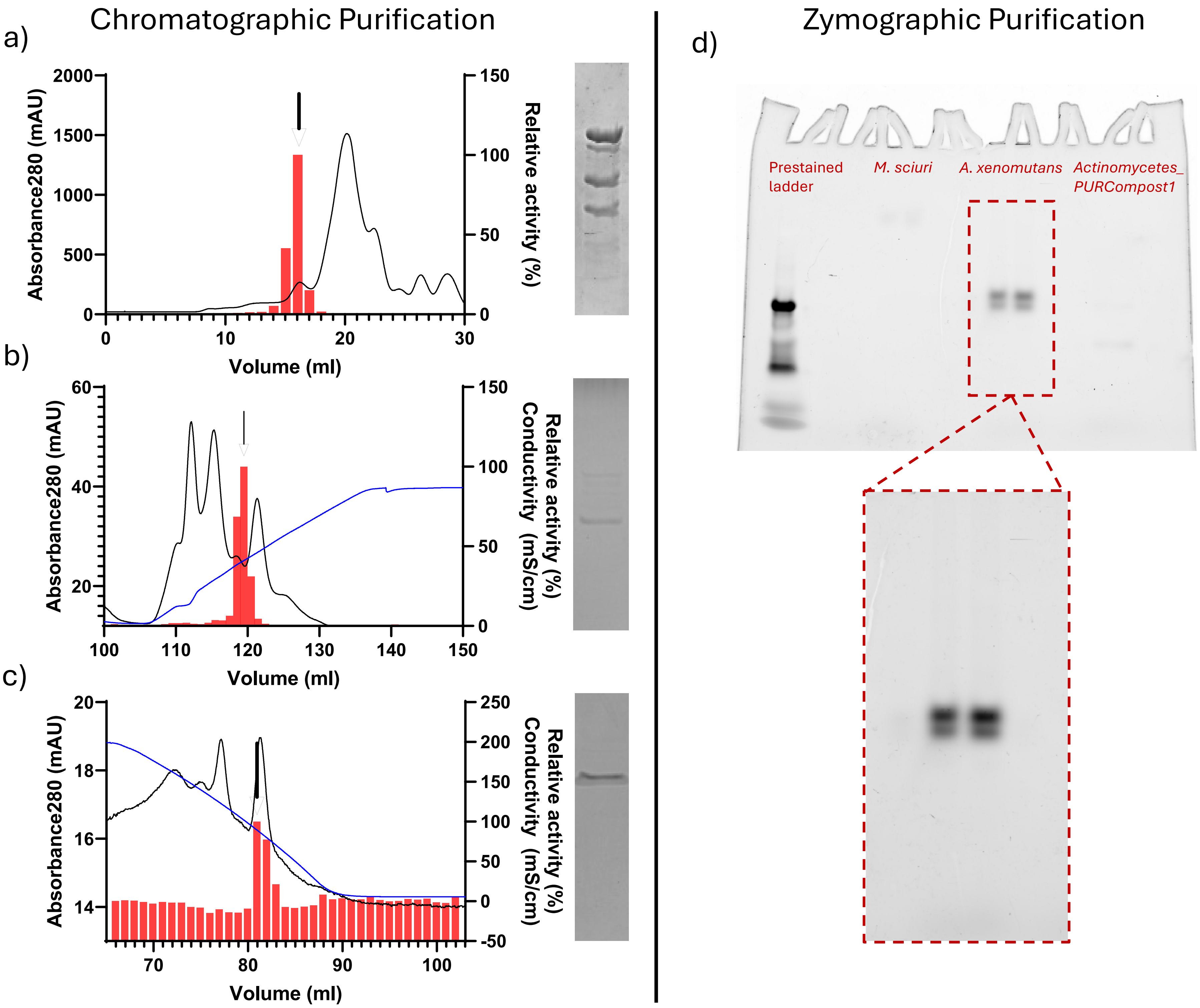


**Supplementary Figure 5. Isolation and purification of plastic degrading enzymes from bacterial isolates**

(a) Size exclusion, (b) hydrophobic interaction, and (c) anion exchange chromatography chromatograms from the purification of *B. licheniformis* overlaid onto the relative activity of each collected fraction on the PURase probe (PURp1), selected fractions chosen for further purification (a and b) or in gel digest (c) are marked with black arrows. A conductivity curve is also present for anion exchange and hydrophobic interaction chromatography. (d) Native zymography gel of anion exchange fractions from *A. xenomutans, M. sciuri,* and Actinomycetes_PURCompost1 post incubation with PURase probe (PURp1). Also, PageRuler™ Plus Prestained Protein Ladder is included in the gel for verification of fluorescence.

### Figure S6

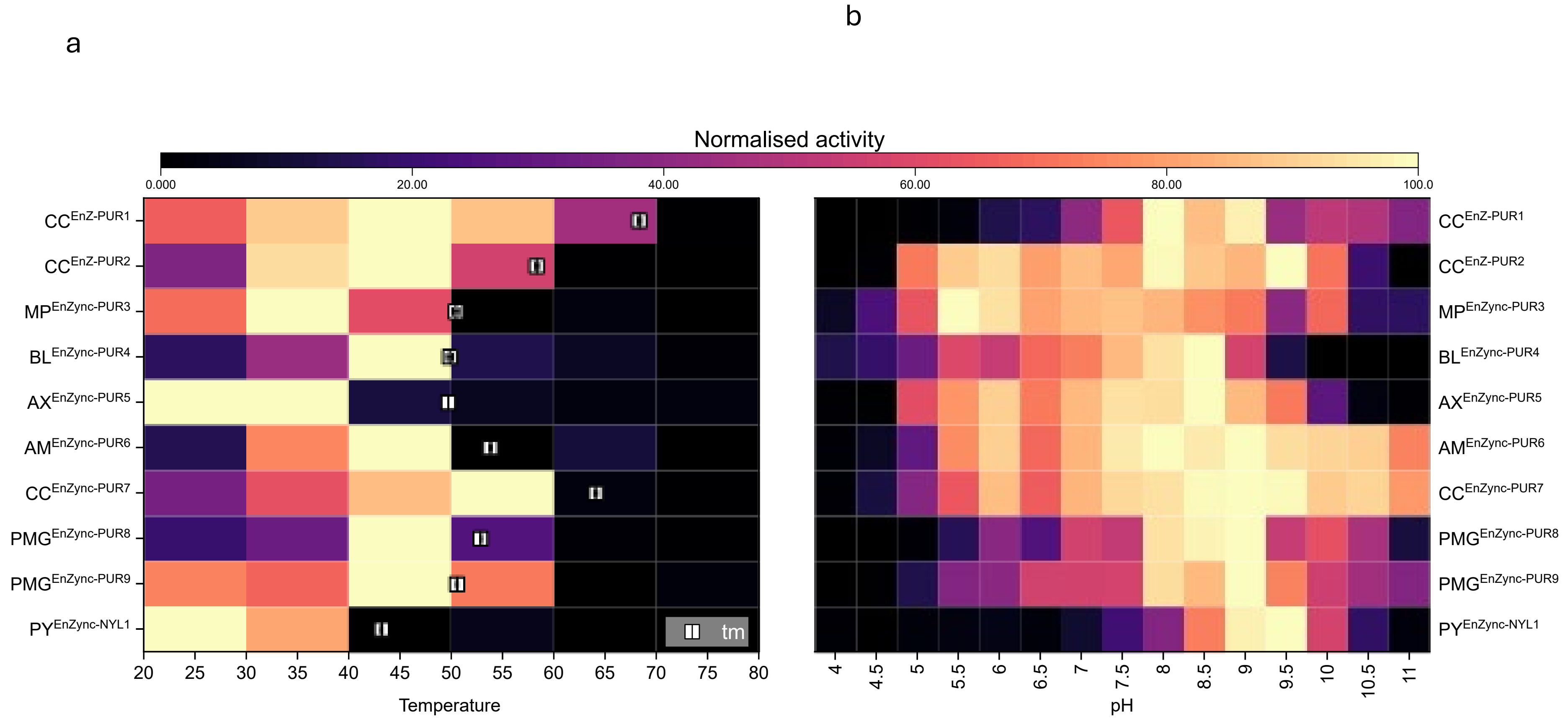


**Supplementary Figure 6. Optimum reaction conditions and melting temperatures of identified enzymes.**
Optimum activity on PURp1. (a) Relative activity as a function of temperature. White squares indicate melting points determined by differential scanning fluorometry (n=3-4). (b) Relative activity as a function of pH. 4-6 in 100mM Na-citrate, 6.5-9.5 100mM Bis-Tris propane, 10-11 in 100mM CAPS. All reactions were performed in the presence of 100mM NaCl.

### Figure S7

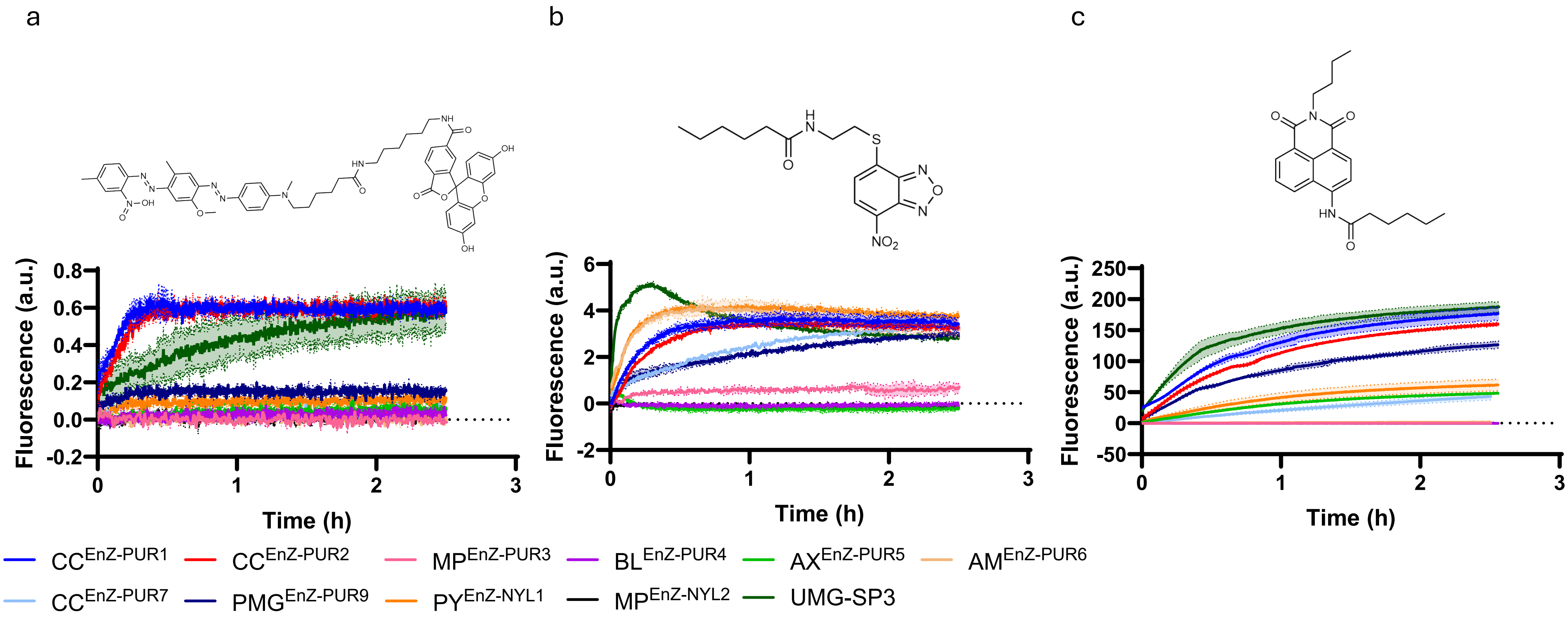


**Supplementary Figure 7. Screening of NYLONase potential with nylonase probes.** Kinetics of cleavage of (a) NYLp1, (b) NYLp2 and (c) NYLp3 (c) in the presence of 1µM of different enzymes. Fluorescence is showed as means and error bars as standard error of means. Control samples containing only the probes have been substracted.

### Figure S8

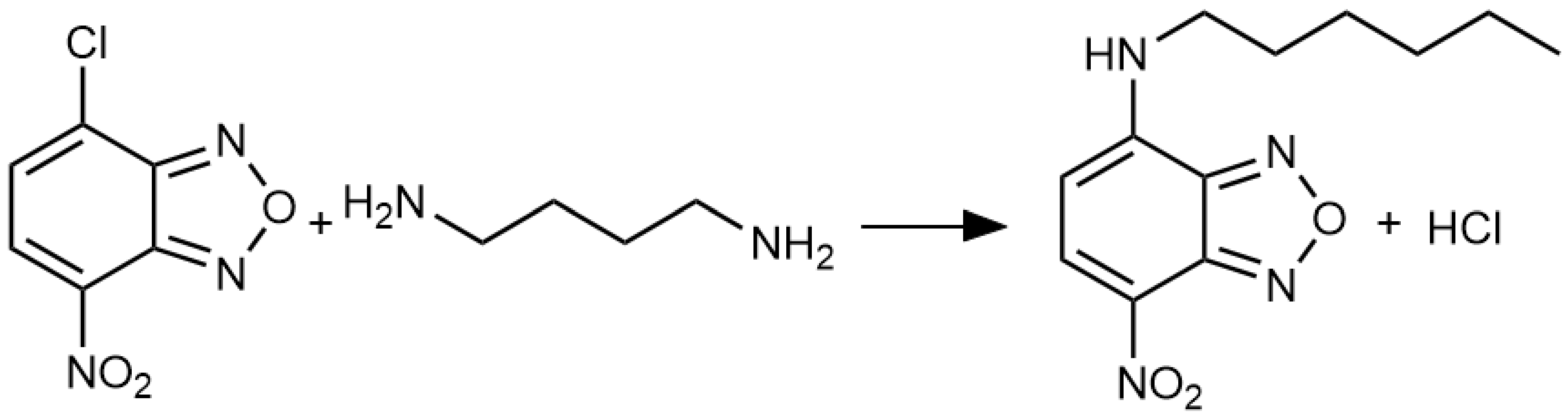


**Supplementary Figure 8.** NBD-Cl reaction in presence of HMDA.

### Figure S9

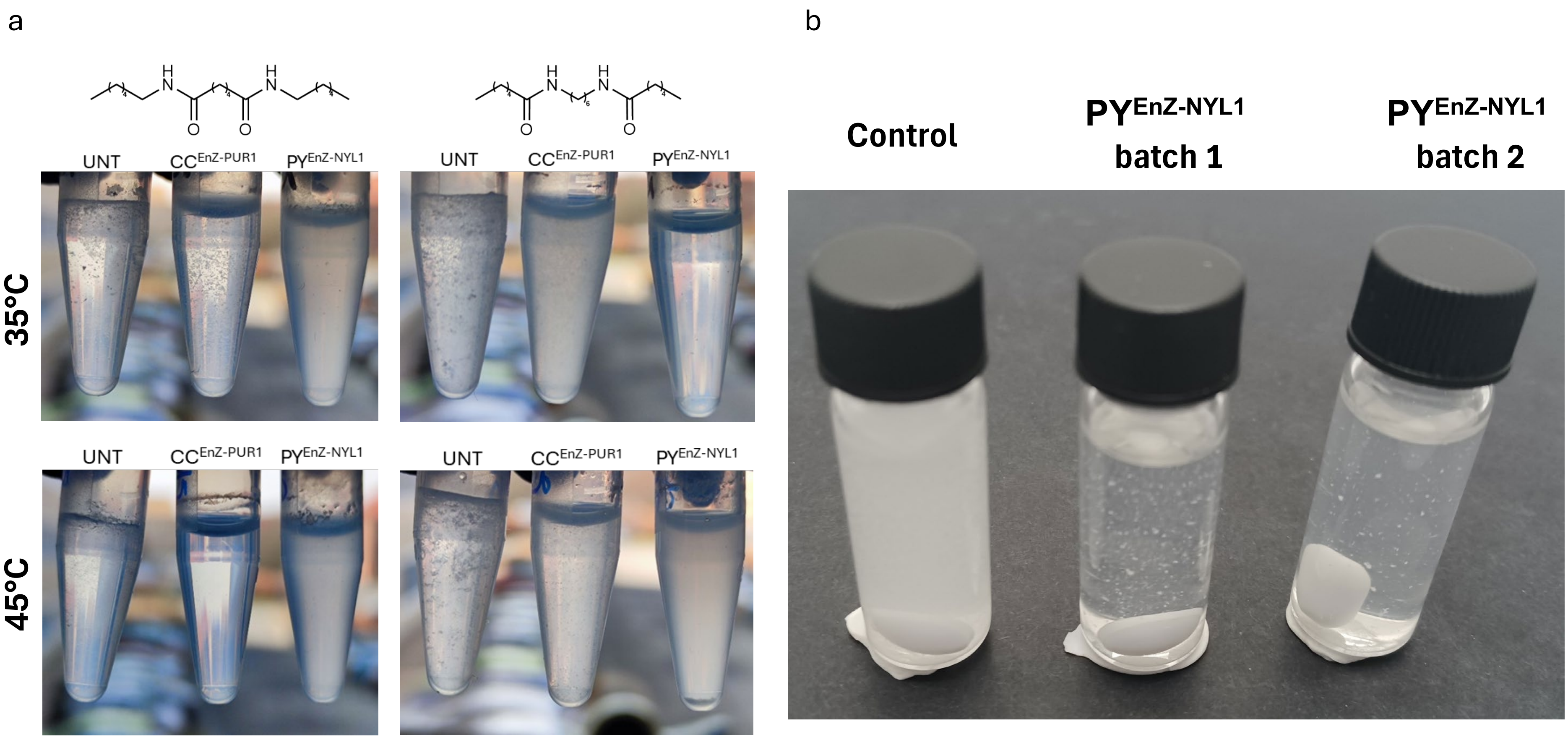


**Supplementary Figure 9. Screening of nylon-degrading potential.** (a) Turbidity after 7 days of incubation of 2 mM nylon dimer samples with 1 μM of CC60 and PY69. (b) Degradation of 2 mM N-mimic after 72 h in the presence or absence of 2 μM of PY69 from two different production batches at room temperature.

### Figure S10

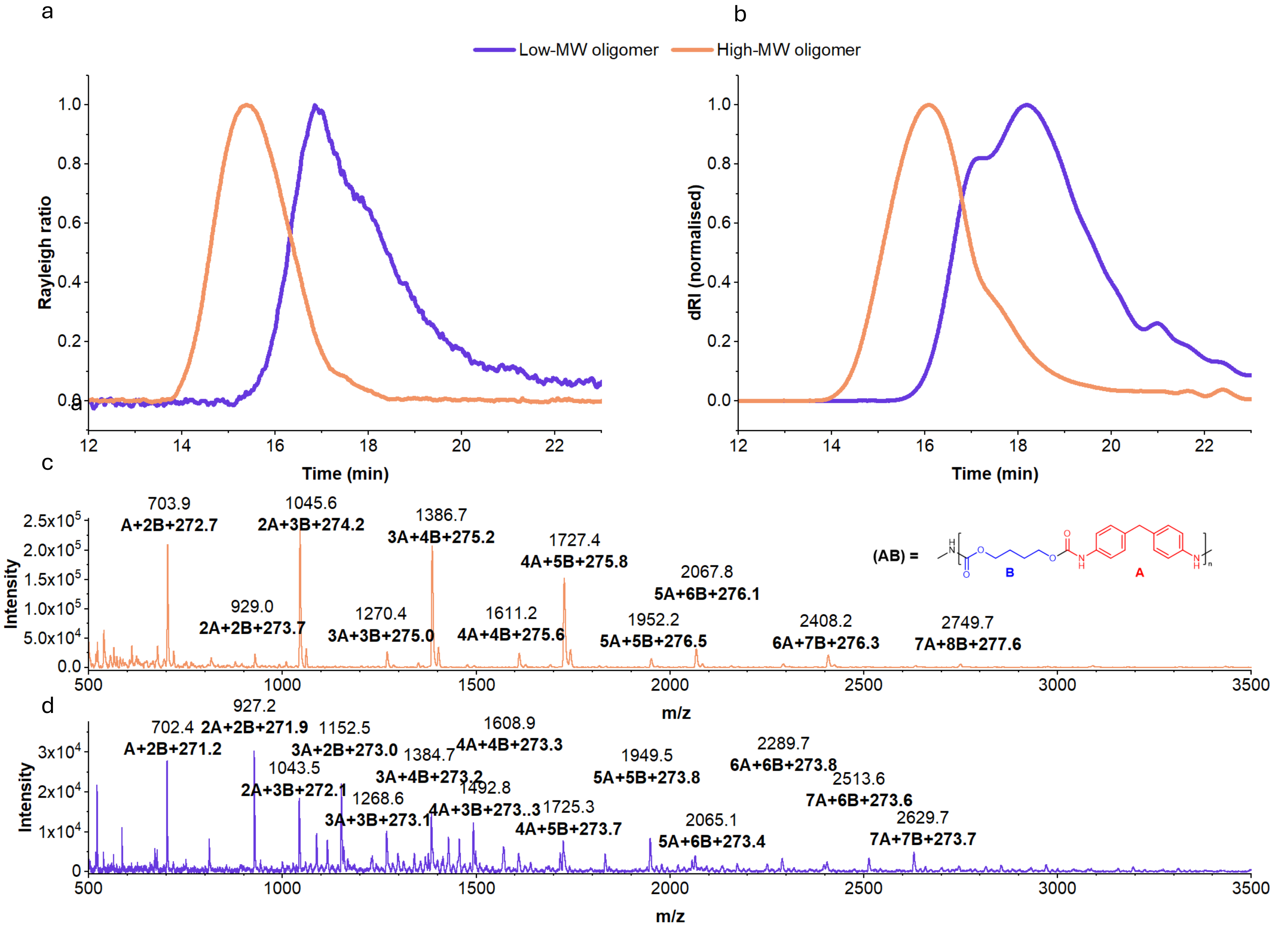


**Supplementary Figure 10. Characterization of PUR oligomers.**
**A, B:** Gel permeation chromatography (GPC) of PUR oligomers dissolved in DMF + LiBr. Showing traces from the light scattering detector of oligomers (A) and refractive index detector (B).
**C, D:** MALDI-TOF mass spectrometry of PUR oligomers. Peak at ~703 is interpreted as BAB+$H^+$ which formally has a mass of 431.2 and an unknown end modification. All interpreted peaks are assigned with their m/z value and their interpretation in bold.
**C)** High-MW oligomer peaks are regularly spaced at 340.1 ± 0.2 Da, consistent with the mass of a single AB segment. Minor peaks spaced 224 Da from the major peaks correspond to MDA groups. An unexplained mass of around 275Da is an unknown modification.
**D)** Low-MW oligomer spectra also show peaks at 340.1 ± 0.2 Da, but dominant peaks are spaced 448 Da apart from the peaks observed in the high-MW oligomer, suggesting end capping with two MDA groups. An unexplained mass of 273 Da is an unknown modification to all peaks.

### Figure S11

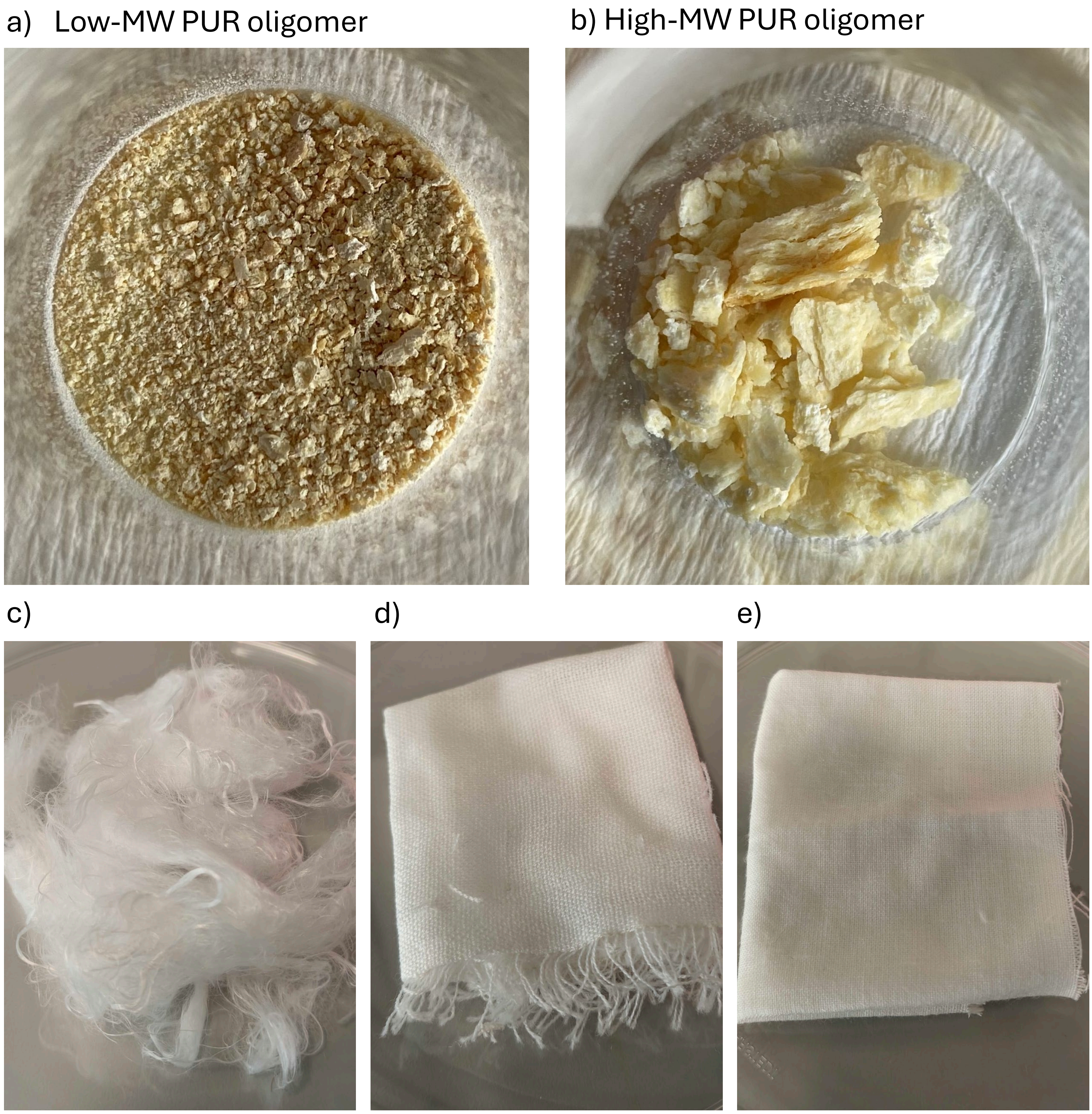


**Supplementary Figure 11. Visual appearance of polyurethane and nylon insoluble samples.** Polyurethane oligomers with (a) low and (b) high molecular weight. Nylon 6,6 (c) threads sample 1 (20-30 µm) and textile (d) sample 2 (30-40 µm) and (e) sample 3 (20-30 µm).
